# Social caste switching triggers emergence of novel cellular identities in *Zootermopsis* termites

**DOI:** 10.64898/2026.04.12.718059

**Authors:** Catherine Gatt, Kensei Kikuchi, Cong Liu, Esra Kaymak, Simon Hellemans, Fuka Koja, Thomas Bourguignon, Fabio Zanini

## Abstract

A hallmark of eusocial insects is the specialisation of individuals into castes that share the same genome but present distinct phenotypes. Caste switching can affect gene expression^1,2^, but can it give rise to entirely new cell types? Here, we generated single-cell transcriptomic atlases of a king, a queen, a soldier, and six workers of the termite *Zootermopsis nevadensis* to understand whether caste differentiation is accompanied by the emergence of novel cellular identities. Out of 24 annotated termite cell types, 18 of which mirrored homologous *Drosophila* cell types, one was unique for king, one was almost entirely restricted to the queen, and none were exclusive to nonreproductives. Instead, nonreproductive termites possessed more muscle and neuronal cells and less fat cells than reproductives. Caste-linked transcriptional signatures were detected in most cell types, including rare genes expressed with dual specificity for both cell type and caste. Despite the emergence of new cell types in reproductives, expression of hormones, immune genes, carbohydrate-active enzymes^3^, and rapidly adapting genes^4^ were only weakly altered by caste. These findings show that cellular differentiation can be induced anew in adult organisms without dysregulating their key physiologic pathways.

## Main

Social insects form colonies of related individuals based on division of labour. A reproductive caste, typically represented by a few individuals, called royalty, is surrounded by many more colony members who perform non-reproductive tasks such as foraging and defense^5^. All colony members form a family unit; the differences in caste phenotype and behaviour are rooted in differential gene expression^6^. Although many genes are differentially expressed among castes in social Hymenoptera^7,8^ and termites^9,10^, the largest non-hymenopteran social insect lineage^11^, they rarely overlap across species^10,12^. An overarching explanation for the molecular basis of insect sociality is missing.

Given the central role of cells for organismal physiology and development^13^, we hypothesised that castes might possess not only unique transcriptional patterns, but entirely new cell types. The differentiation program for such cells would remain cryptically encoded in the genome throughout adulthood and only manifest itself upon caste switching, driven by social and biochemical cues. Can caste differentiation really generate new cellular identities?

Here, we report the discovery of caste-restricted cell types in a termite. Building upon work on ant and bee brains^14–17^, we construct a whole-body transcriptomic cell atlas of the termite *Zootermopsis nevadensis*. We used three castes: totipotent active immatures (workers), a sterile worker-derived soldier, and a pair of unwinged reproductives (queen and king) that lost their wings before founding a new colony^18,19^. All cells found in non-reproductives were also found in royalty, while the king and queen each also possessed a single unique cell type. This pair of cell types was accompanied by differential cell type abundance and transcriptional plasticity within shared cell types, including neurons, muscle, fat, and stem cells. Despite such sweeping changes, no caste preference was found at the cell type level in the expression of developmental hormones, immune genes, carbohydrate-active enzymes^20^, and rapidly adapting genes, indicating that the activation of new cell differentiation programs can coexist with stable homeostasis across many physiological axes. This study demonstrates the existence of latent cell differentiation programs in adult organisms and traces a path towards a deeper understanding and engineering of such programs in other organisms^21^.

### A single-nuclei transcriptomic atlas of the termite *Zootermopsis nevadensis*

To characterise the cellular landscape of a social insect, we focused on *Zootermopsis nevadensis*, which has a high-quality genome assembly^4^ and can be bred in a laboratory. We developed an experimental workflow, which combines gentle dissociation, fluorescence-activated nuclei sorting, droplet-based libraries, and short-read sequencing, to obtain high-quality single-nuclei transcriptomic data from whole individual termites (**Figure 1A** and see Methods). We processed one library per caste and obtained 2,288 nuclei from one king, 6,529 nuclei from one queen, 4,805 nuclei from one soldier, and 10,630 nuclei from a pool of six individuals of the worker caste, for a total of 24,252 nuclei and 13,413 genes (**Figure 1A**).

**Figure 1:**
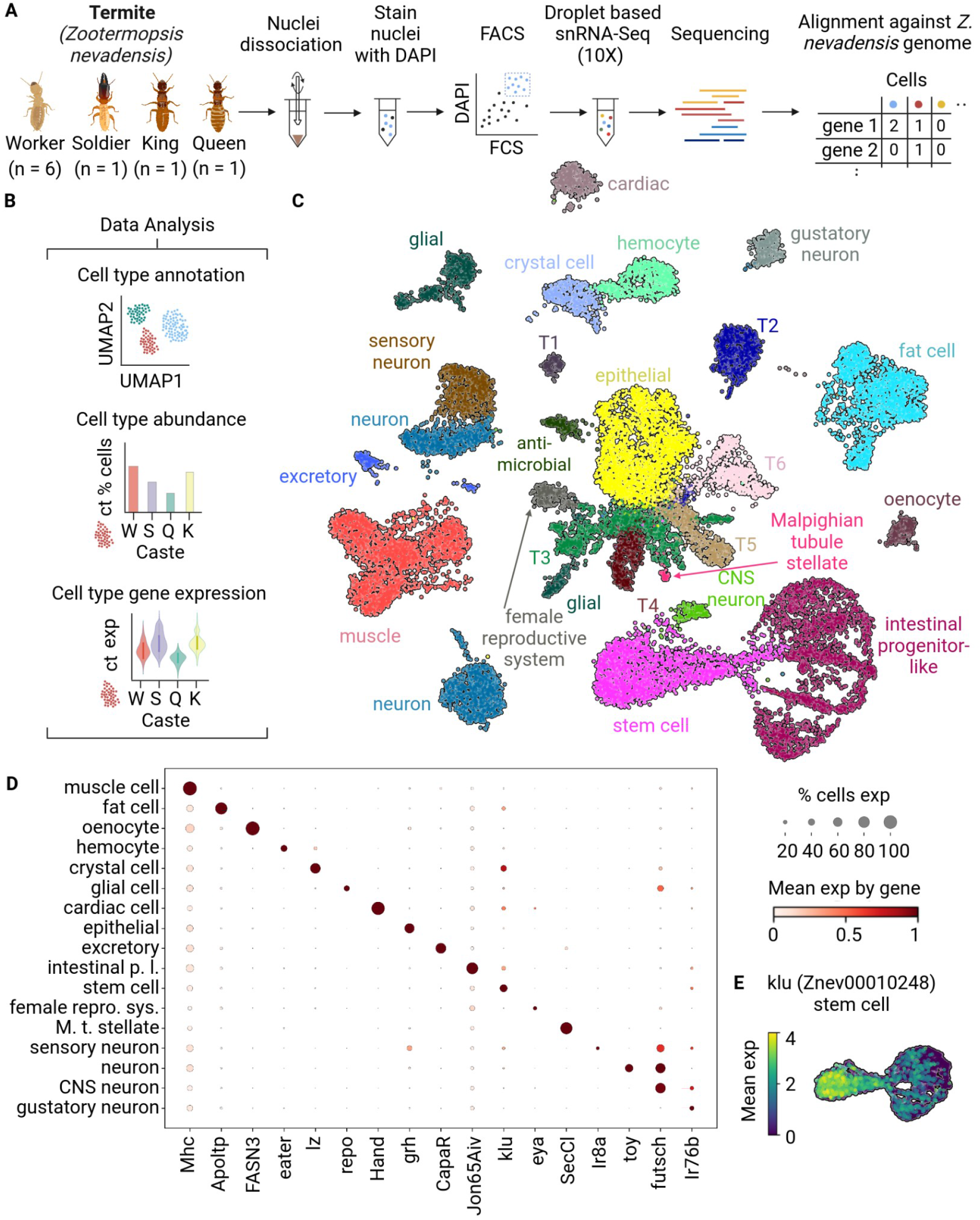
Whole-body cell atlas of the termite *Zootermopsis nevadensis*. **A)** Schematic of the single-nuclei dissociation, sequencing, and cell type annotation workflow. **B)** Palette of computational analyses performed on the *Zootermopsis* atlas data (W = Worker, S = Soldier, Q = Queen, K = King). **C)** UMAP embedding of all nuclei coloured by cell type as assigned by comparison against *Drosophila melanogaster*. Cell types with no detectable homology were labelled T1-T6. **D)** Dot plot showing the expression of termite homologs of *Drosophila* cell type markers across termite cell types. Dot size represents the proportion of cells expressing the gene and colour (white to dark red) shows average expression in that cell type. **E)** UMAP embedding of stem cell and intestinal progenitor-like cells, coloured by expression of the *Zootermopsis* homolog of *Drosophila* stem cell marker *klu*.

Single-nuclei data from all castes were preprocessed and clustered into 33 cell populations of unknown identity (see Methods, **Supplementary Figure 1B**). To annotate these clusters, cross-species data harmonisation against the cell atlas of *Drosophila melanogaster*^*22*^, which diverged from termites 400 million years ago^23^, was used, combining the graph-based algorithm SAMap^24^, the deep neural network model SATURN^25^, the recent SATURN-xfer^26^, and manual refinement and merging based on orthologous marker genes. Overall, 24 cell types were annotated and embedded with no need for caste harmonisation^27^ (**Figure 1C**). Of those, eighteen cell types could be confidently named and expressed canonical marker genes (**Figure 1D**) and six cell types (T1-T6), each expressing individual marker genes (**Supplementary Figure 1D**), were treated as novel. Further transcriptional heterogeneity was detected within some clusters, including stem cells, which presented a decreasing gradient of the conserved marker *klu* towards intestinal progenitor-like cells (**Figure 1E**). Overall, these findings demonstrated robust acquisition and annotation of single-nuclei data from non-model insects despite the scarcity of experimental tools and a large evolutionary distance from the closest annotated species.

### The abundance and transcriptional identity of cell types are caste-specific

*Zootermopsis* is characterised by a working stage that can optionally differentiate further into soldiers or reproductives - king or queen (**Figure 2A**). To understand how caste affects cellular identities, we compared the relative abundance and transcriptional profiles of each cell type across these castes.

**Figure 2:**
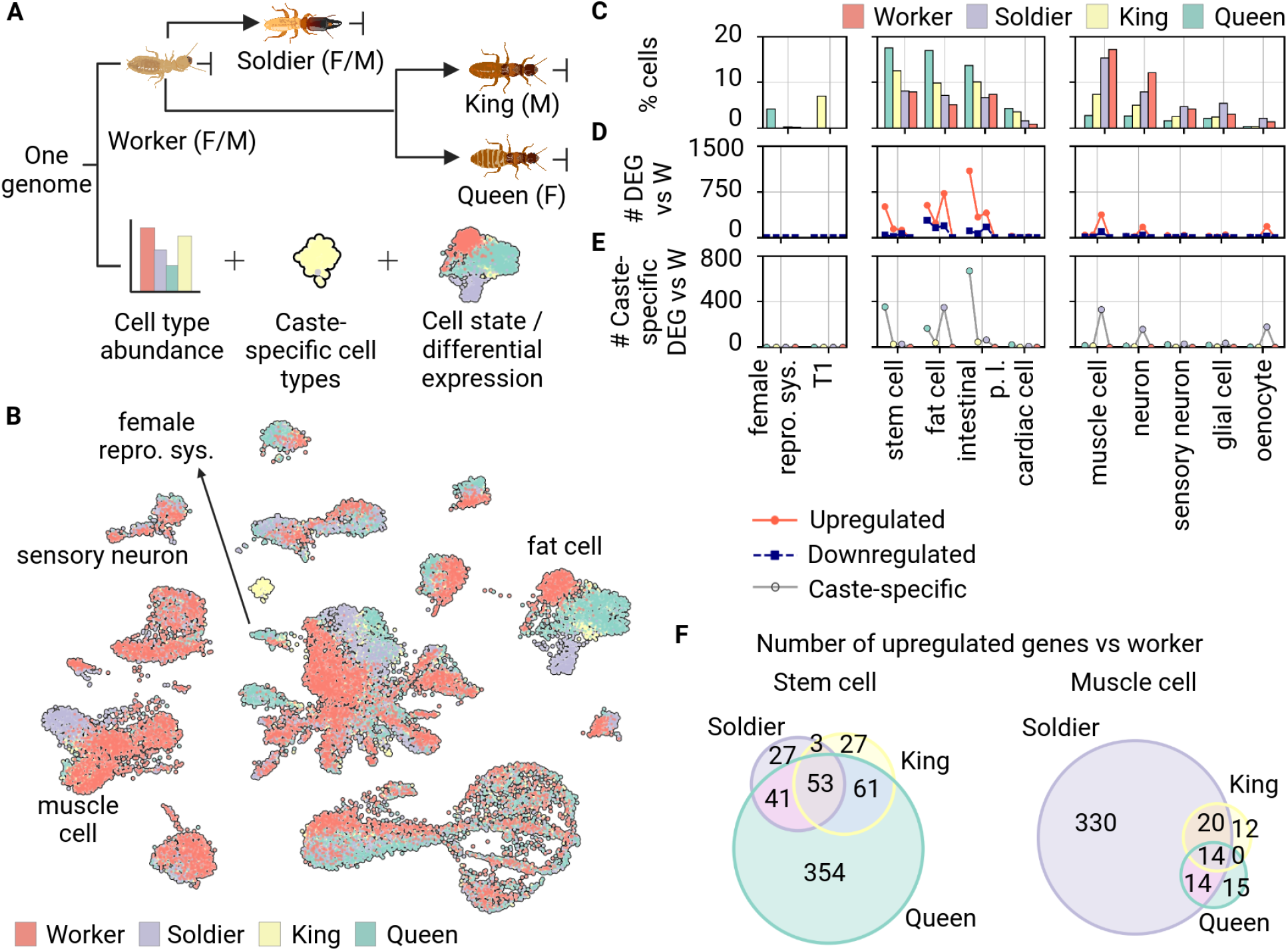
Emergence of new cell types in termite reproductives. **A)** Diagram of developmental and caste transitions in the termite *Zootermopsis nevadensis* and of the different analyses comparing castes at the single-cell level. **B)** UMAP embedding of termite nuclei as in **Figure 1C**, coloured by caste of origin. **C)** Relative cell type abundance across castes, including caste-specific (left) and shared (right) cell types. **D)** Number of upregulated and downregulated genes in each caste compared to workers (p-value < 0.05), for each cell type. **E)** Number of genes exclusively upregulated in this one caste compared to workers (p-value <0.05), for each cell type. **F)** Venn diagram of upregulated genes against workers in queen, king, and soldier in stem (left) and muscle cells (right).

Cell embeddings coloured by caste of origin provided a first indication that cell types might not be evenly distributed amongst castes (**Figure 2B**). To quantify this observation, we computed the relative abundance of each cell type and caste over all cells from the same caste (**Figure 2C**). Two cell types were highly specific to one caste. Female reproductive cells constituted 7% of sampled nuclei in queen, less than 1% in soldier and the working castes - in agreement with morphological observations of termite gonads^28^, and were absent in the king termite. T1 cells, for which no sister cell type could be safely identified in *Drosophila*, were found exclusively in *Zootermopsis* king (**Figure 2C**). Multiple genes upregulated in T1 versus other cells might be related to lipid transport and immunity (**Supplementary Table 5**). Nonreproductive castes lacked exclusive cell types. Among other cell types, stem, fat, intestinal progenitor, and cardiac cells were more abundant in termite reproductives, whereas muscle, neurons, glial cells, and oenocytes were more abundant in nonreproductive castes (**Figure 2C**).

Given that queens, kings, and soldiers all derive developmentally from workers, we asked how many genes are upregulated in cells from each derived caste versus worker cells of the same type. We found that cell types that were more abundant in reproductive castes (e.g., stem cells) often also upregulated many genes compared to worker cells of the same type (**Figure 2D**), suggesting that increased cellular abundance is accompanied by activation of new transcriptional programs. We then asked whether this expanded transcriptional footprint is shared across non-worker castes and found a highly dynamic landscape across cell types (**Figure 2E**). In some cell types, most upregulated genes were exclusive to one caste (e.g., soldier for muscle cells), whereas other cell types upregulated common genes during multiple caste transitions (e.g., for fat cells). These data demonstrate that caste transitions cause bidirectional changes in cell type abundance, transcriptional tuning of extant cell types, and emergence in reproductives of novel cell types that might or might not be already present in alates, the individuals that eventually become royalty after a nuptial flight.

### Interaction of cell type and caste leads to dual genetic specialisation

The observed changes upon caste differentiation raised the hypothesis of dual genetic specialisation, i.e., transcripts expressed only in one specific combination of caste and cell type. To test this hypothesis, we further examined fat and muscle cells, which exhibited poor overlap between castes in the embeddings (**Figure 3A**). Evolutionarily conserved cell type markers, such as *Apoltp* for fat cells and *Mhc* for muscle, were expressed by virtually all cells within each type regardless of caste (**Figure 3B**). However, we also identified genes with *preferential* dual specialisation, characterised by unique expression by one cell type and, within that type, upregulation in a subset of castes. For instance, *TpnC47D*, a homolog of the human gene *TNNC2*, involved in fast-twitch muscle contraction, was almost exclusively expressed by soldier muscle but not by muscle in other castes or by non-muscle cells from any caste (**Figures 3B-C**). Similarly, almost only fat cells from reproductive castes expressed the *Zootermopsis* homolog of the uncharacterised *Drosophila* gene CG34370 (**Figures 3B-C**).

**Figure 3:**
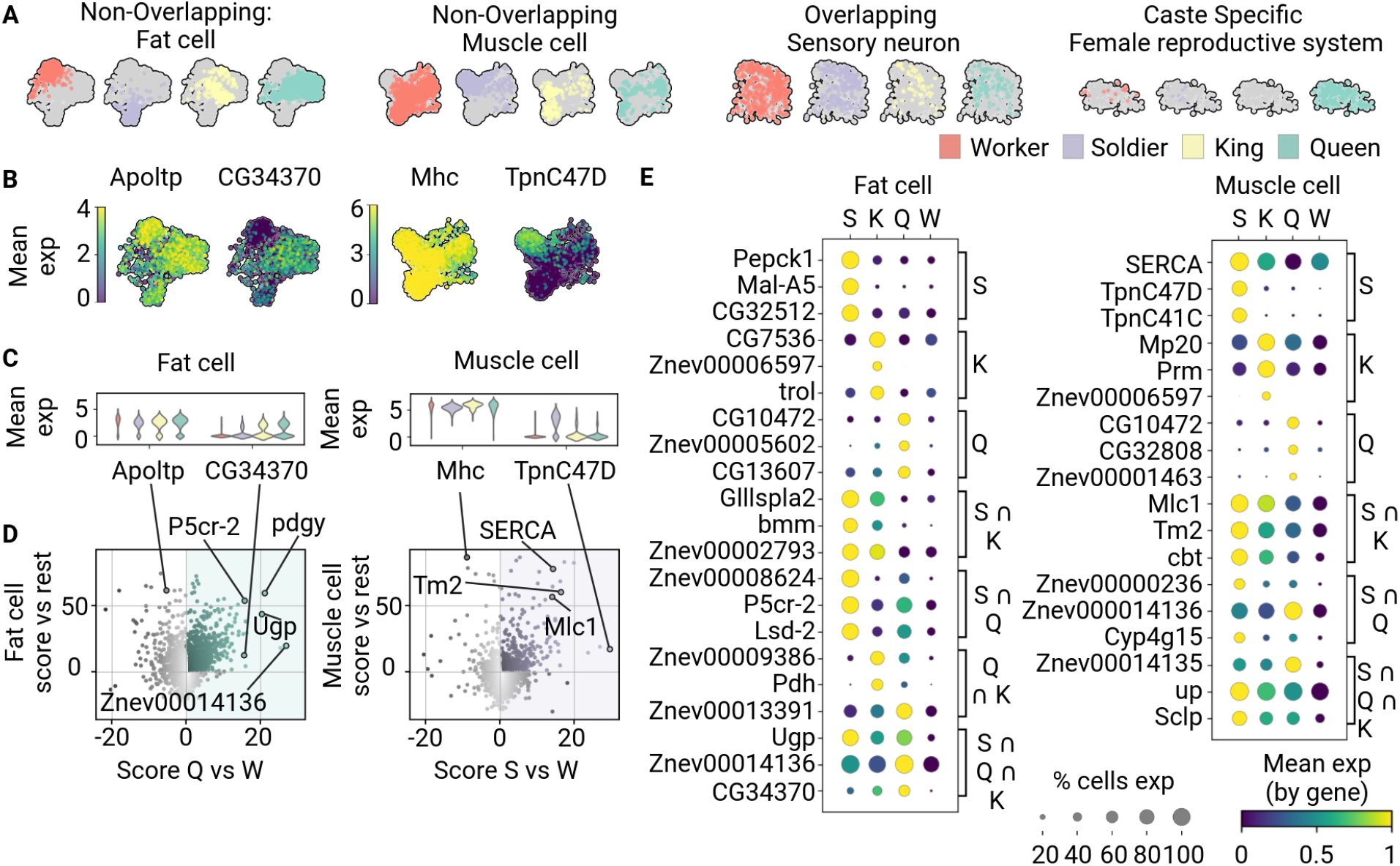
Dual genetic specialisation to cell type and caste. **A)** UMAP embedding for fat cells, muscle, sensory neurons, and female reproductive system cells, for individual castes, showcasing different degrees of overlap between castes within individual cell types. **B)** UMAP embeddings of fat (left two charts) and muscle cells (right two panels) color coded by expression of individual genes. **C)** Violin plots showing the distribution of expression for specific genes in fat (left chart) and muscle cells (right chart) across castes. **D)** Scatter plots for dual differential expression analysis. For each gene, represented by a dot, the x coordinate shows the statistical test score of queen (Q) fat (left) / soldier (S) muscle (right) cells against worker (W) cells of the same type. The y coordinate shows the statistical score of cells of that type versus other cells across all samples. Genes in the top right section are dual specialised. **E)** Dot plots showing dual specialisation genes in fat (left) and muscle cells (right) (W = Worker, S = Soldier, Q = Queen, K = King).

How prevalent are these patterns across the transcriptome? To answer this question, we computed differential expression across two independent axes for each gene (**Figure 3D**): the x-axis represented one caste against the rest within a cell type, while the y-axis cell type against others within a caste. This approach uncovered a highly diverse transcriptional landscape, which included genes that were only specific for cell type but not caste (including *Alpoltp* and *Mhc*), genes that were specific for caste but not cell type, as well as dual specialisation genes (**Figure 3D** and **Supplementary Figure S4**). Many genes exhibited some degree of specificity for either caste, cell type, or both, precluding an exact distinction between these categories. Moreover, some genes with dual specialisation, including CG34370, were not restricted to a single caste (**Figure 3E**). Evidently, in termites, both individual transcripts and broad (transcriptional) cellular identities can dual-specialise for both cell type and caste, a phenomenon not observable in caste-less organisms such as *Drosophila*.

### Cell type and caste jointly determine hormone, immune, and CAZyme usage

Bulk transcriptomics of termite tissues has been used to identify caste-specific differentially expressed genes (DEGs) within important physiological pathways including hormones, immunity, and lignocellulose digestion^2^. What cell types are responsible for those patterns?

Genes in the Juvenile hormone (JH) biosynthesis pathway, associated to moulting and caste differentiation^2,29,30^ showed clear cell type specificities (**Figure 4A**) and were expressed at significantly higher levels in matching cell types in the queen versus the worker (**Supplementary Table S7**). On the contrary, the presumed JH receptor Znev00002391, homolog of *gce*^*31*^, was broadly expressed by many cell types (**Figure 4A**). Expression differences of *gce* between workers and queens or kings were not significant, whereas expression was slightly higher in workers compared with soldiers (**Supplementary Table S8**). These findings support the hypothesis that endogenous JH production is tightly regulated across the termite body while the response to JH stimulation - including exogenous JH - might be widespread across cells and tissues.

**Figure 4:**
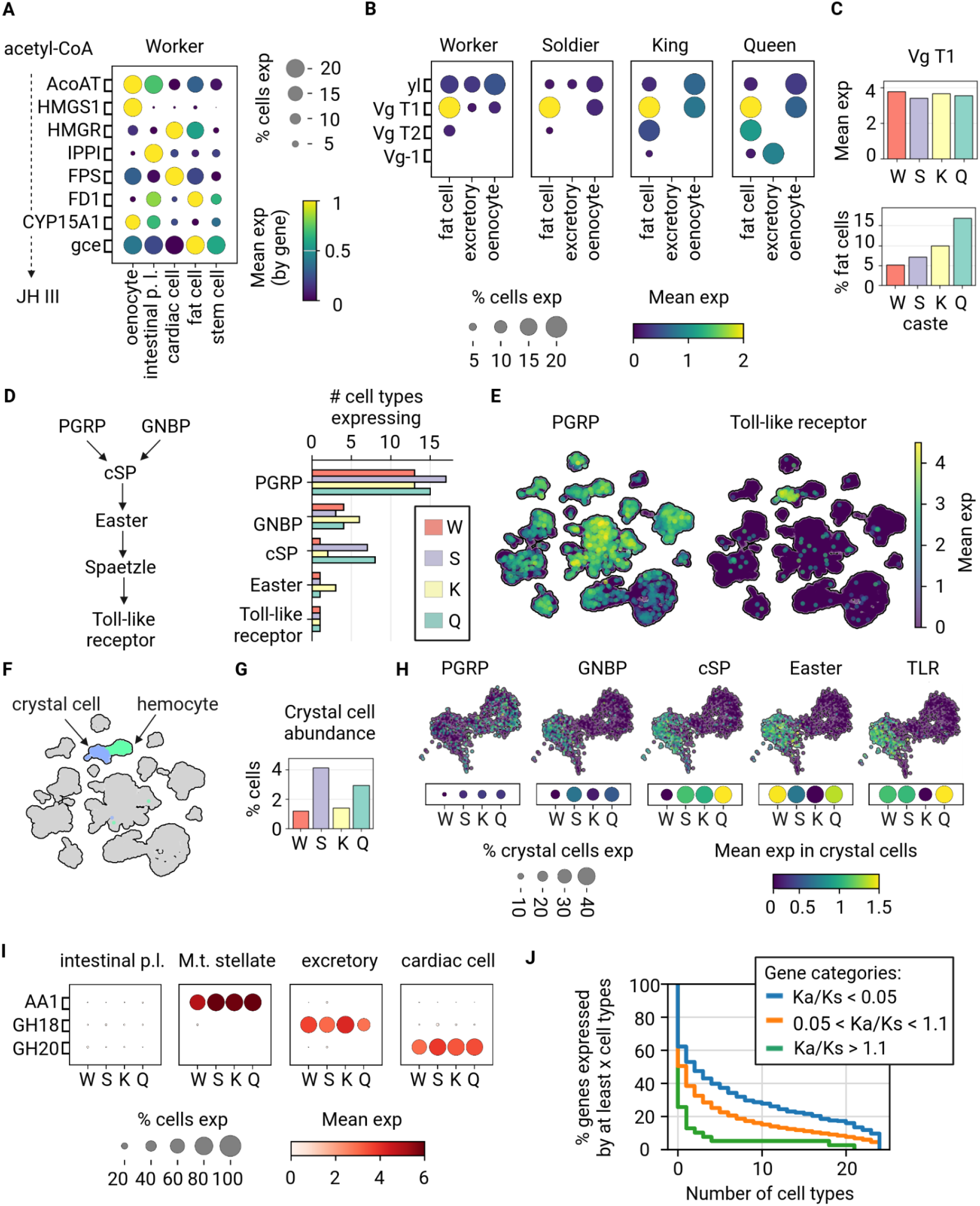
Expression patterns of hormonal, immune, carbohydrate-active enzymes, and rapidly adapting genes. **A-B)** Dot plots showing expression of pathway-specific genes across cell types (columns) and castes (rectangles). The legend is shared between both panels. **A)** Expression of Juvenile Hormone (*JH*) biosynthesis-related genes in the worker and **B)** Vitellogenin (*Vg*)-related genes across all castes. **C**) Bar plots of cell type abundance and transcriptional abundance within the same cell type for *Vg* in fat cells. **D**) Bar plot of the number of cell types expressing genes involved in the toll signalling cascade. **E)** UMAP embedding of all nuclei coloured by expression of PGRP and Toll-like receptor. **F)** UMAP embedding of all nuclei coloured by immune-related cell types, hemocyte and crystal cell. **G**) Bar plot of cell type abundance of crystal cells. **H)** UMAP embedding of crystal cell and hemocyte coloured by expression of genes involved in the toll signalling cascade. **I)** Dot plot showing 3 highly expressed CAZymes transcripts across selected cell types and castes. **J)** Cumulative distribution of the number of cell types in which genes are expressed (mean log expression ≥ 0.1 in ≥ 10% of cells), grouped by low (< 0.05, blue), middle (0.05-1.1, orange) and high (> 1.1, green) Ka/Ks values.

Another gene associated with caste is Vitellogenin (*Vg*), a yolk protein with a proposed role in lifespan extension previously detected at increased levels in queen tissues^2,32^. In our data, the *Zootermopsis* homolog of *Vg, Vg-T1*, was expressed almost exclusively by fat cells, while other homologs had lower expression overall (**Figure 4B**). Remarkably, *Vg-T1* expression in fat cells was comparable across castes, while fat cell abundance was much higher in queen than in other castes (**Figure 4C**). The logical consequence is that physiological changes downstream of *Vg*, including longevity, might be achieved via cell number regulation rather than direct gene regulation.

Caste development has been implicated in termite immunity^33^. Insects rely on pathogen-associated molecular pattern (PAMP) recognition via peptidoglycan-recognition proteins (PGRPs) and the gram-negative binding proteins (GNPBs), which funnel information through intracellular and paracrine signalling networks, including Toll-like receptors^34^. In our data, PAMP recognition genes were broadly expressed but genes encoding proteins deeper in the signaling cascade were expressed in fewer cell types (**Figure 4D**). The only Toll-like receptor expressors were crystal cells (**Figure 4E**), a specialised immune population related to hemocytes (**Figure 4F**). Crystal cells represented between 1 and 4% of all nuclei across castes (**Figure 4G**) and co-expressed many immune genes in the same pathway (**Figure 4G**). Expression of individual genes varied across castes, suggesting that the broad mechanisms of immunity are shared across castes but not the exact expression levels (**Figure 4H**).

Endogenous carbohydrate-active enzymes (CAZymes) have been hypothesised to assist gut microbes during lignocellulose digestion, an ecologically central function of termites^20,35,36^. However, in our data, CAZymes were not expressed by intestinal (progenitor) cells. Instead, multiple CAZymes were expressed specifically in some cell types: AA1 (laccase) in stellate, GH18 (chitinase) in excretory, and GH20 (β-N-acetylhexosaminidas) in cardiac cells (**Figure 4I**). For each cell type considered, caste of origin did not seem to affect CAZyme expression (**Supplementary Figure S9**). These results indicate that over evolutionary time, individual CAZymes were integrated into preexisting gene regulatory networks unrelated to lignocellulose processing.

To test across the entire transcriptome whether genes under positive selection tend to associate with cell type- and/or caste-specific expression, we analysed the ratio of nonsynonymous to synonymous changes across termites for all genes (Ka/Ks)^37^. We found that genes under putative positive or relaxed selection (Ka/Ks > 1.1) were expressed by significantly fewer cell types than genes under purifying selection (Ka/Ks < 0.05), indicating that selection acts more strongly on widely used genes (**Figure 4J**).

## Discussion

In *Zootermopsis*, two cell types were restricted to the reproductive caste and none to nonreproductive castes. This discovery raises the immediate question as to whether vertebrate eusocial organisms such as the mole rats^21^ induce novel cell differentiation in adults upon social organisation. If so, the possibility of engineering such novel cells in caste-less organisms would appear less remote than initially thought.

Many cell types showed expression of canonical cell-type markers across castes but also of caste-specific genes. For example, soldier-specific expression in muscle cells of genes associated with calcium signalling and fast-twitch fibre contraction might relate to biting/crushing behaviours used by *Zootermopsis* soldiers during combat with enemies^38^. In addition to new cell types and adaptation of extant ones, a third mechanism of caste adaptation was observed: cell number regulation. For instance, the *Zootermopsis* homolog *Vg-T1* of vitellogenin, previously associated with queen development and ageing^39^, was specifically expressed by fat cells at similar levels across castes. However, the relative number of fat cells was higher in the queen compared to other castes, leading to higher expression *across the whole body* in queens.

Single-nuclei isolation required only individual specimens (queen, king, soldier) or a small pool (six workers). Thanks to recent algorithmic advances^24–26^, cell types could be annotated robustly based on data from *Drosophila*^*22*^, which diverged from termites about 400 million years ago^23^. This combination of highly sensitive experimental technologies and computational tools appears poised to enable cell atlases from single-digit available specimens of rare, uncharacterised invertebrates at planetary scale.

In summary, we found direct evidence that caste specialisation involves (i) forging novel cell types, (ii) altering gene expression (*i*.*e*., “cell state”) within existing cell types, (iii) adjusting the relative abundance of cell types, and (iv) restricting expression of genes undergoing rapid changes to select cell types. These findings paint a multiscale picture of caste development connecting individual genes to cells, organisms, and entire insect societies.

## Data Availability

The sc-RNA seq FASTQ files, count barcodes, matrices and features from each caste, and gene cell count object with processed reads, annotated cell types and *D. melanogaster* orthologs as described above are available on the Gene Expression Omnibus (accession no. GSE323970).

## Code Availability

The code used for this manuscript is publicly available at https://github.com/fabilab/zootermopsis_atlas_paper. An archived version is available on Zenodo at the following DOI: https://doi.org/10.5281/zenodo.18946121.

## Contributions

F.Z. and T.B. conceptualised the study. K.K. collected the samples. C.G., E.K. and F. K. performed lab experiments and generated data. C.G., C.L., S.H. and F.Z. analysed the data. C.G. and F.Z. wrote the original draft manuscript. All authors edited the manuscript.

## Methods

### Colony Foundation

A *Z. nevadensis* colony was collected from pine forests in Kawanishi City, Hyogo Prefecture, Japan, in May 2023. The colony was maintained at 27°C and 80% relative humidity under constant darkness until it produced nymphs that matured into alates and started to swarm about one month after the colony was collected. Newly emerged alates were separated into sex-specific plastic boxes containing moistened filter paper and male alates were paired with female alates. Each pair was then transferred to a 15 ml conical tube filled with pine dust and cellulose powder and maintained at 27°C and 80% relative humidity under constant darkness. Each pair founded an incipient colony composed of one queen, one king, and their first broods, which included a dozen workers and one soldier at the time specimens were collected for nuclei dissociation. Of note, the incipient colonies used in this study consisted of alates originating from a single mother colony that were reared under identical conditions.

### Sample Preparation

Four different types of samples were collected from these incipient colonies for single-cell sequencing. Six workers were selected from a single incipient colony on October 25, 2023; one queen and one soldier were selected from a single incipient colony on November 1, 2023; one king was selected from a single incipient colony on November 22, 2023. Selected specimens were processed over two successive days. Nuclei dissociation, FACS sorting, and all single-cell sequencing steps until cDNA quantification (end of Step 2.4 Chromium Next GEM Single Cell 3’ Reagent Kits v3.1 (Dual Index) CG000315 Rev E) were performed the same day specimens were selected, as described below. Library construction (Step 3 Chromium Next GEM Single Cell 3’ Reagent Kits v3.1 (Dual Index) CG000315 Rev E) was performed on the following day, as described below.

### Nuclei dissociation

Dissociation and isolation of single nuclei from the whole body of each sample was performed using an adapted version of a protocol established for *Drosophila*^*40*^. Briefly, 100 μl of homogenisation buffer was dispensed into a DNA LoBind 1.5 ml tube (Eppendorf 022431021) and placed on ice. The homogenisation buffer consisted of nuclease free water, 250 mM nuclease free sucrose (FUJIFILM Wako Pure Chemical Industries, Ltd. 19-0001), 10 mM Tris (pH 8.0) (Nippon Gene 312-90061), 25 mM KCl (Cayman Chemical Company 600213), 5 mM MgCl2 (Serva 39772), 0.1% Triton-x 100 (FUJIFILM Wako Pure Chemical Industries, Ltd. 168-11805), 0.5% RNasin Plus (Promega N261B), 1× protease inhibitor (Promega G6521), and 0.1 mM DTT (Formedium Ltd. DTT005). Samples were placed into the homogenisation buffer and nuclei were released by approximately 20 gentle strokes of a plastic pestle (Violamo 1-2955-01) and an additional 900 μl of homogenisation buffer was added to the tube. Each sample was centrifuged (Eppendorf 5424R) at 1000 g for 10 minutes at 4°C, the supernatant was discarded, and the nuclei were resuspended in 1000 μl of resuspension buffer on ice. The resuspension buffer consisted of 1× PBS (pH 7.4) (Thermo Fisher 2669978), 0.5% BSA (Wako Pure Chemical Industries, Ltd. 017-21273), and 0.5% RNasin Plus (Promega N261B). Each sample was then passed twice through a 100 µm cell strainer (As One 100-98-5) and placed into a 5 ml FACS tube (Falcon 352235) with a 35 µm cap for FACS sorting.

### FACS sorting

Following dissociation, nuclei were stained with 1 μl of DAPI (Invitrogen D1306) and were isolated from remaining debris with Fluorescence Activated Cell Sorting (FACS) using a BD FACSAria III. The worker sample was sorted into a 1.5 ml tube containing 400 μl of resuspension buffer. The queen, soldier, and king samples were sorted into 1.5 ml tubes containing 200 μl of resuspension buffer. Immediately after FACS sorting, nuclei were centrifuged at 1000 g for 10 minutes at 4°C and the 700 μl of supernatant was discarded. Nuclei were resuspended in the remaining 50-60 μl of supernatant, and the concentration of nuclei was calculated using a Thoma cell counting plate (Watson 177312C) with a ZEISS Axio Scope A.1 microscope.

### Single-nuclei library preparation

Sequencing was performed as per the 10x Genomics User Guide (CG000315 Rev E). Gel Beads-in-emulsion (GEMs) were generated using the Chromium X Instrument (10x Genomics, Pleasanton, CA, USA), and scRNA-seq libraries were constructed using the Chromium Next GEM Single Cell 3’ Reagent Kit v3.1 (Dual Index). The worker sample was loaded in the Chromium Next GEM Chip G with a target output of 10,000 nuclei, and the queen, soldier, and king samples were loaded in the Chromium Next GEM Chip G with a target output of 5,000 nuclei. All reactions were performed in the Eppendorf Mastercycler X50s with a 96-Deep Well Reaction Module. For both cDNA amplification and sample index PCR, 14 cycles were used across all samples.

### Sequencing

Sequencing was performed on an Illumina NovaSeq 6000 Sequencing System (Illumina, San Diego, CA, USA) using an SP flow cell with paired-end and dual indexing, set for a total of 138 cycles (28 cycles for Read 1, 10 cycles for the i7 index, 10 cycles for the i5 index, and 90 cycles for Read 2).

### Data preprocessing

A custom reference genome of *Z. nevadensis* was generated using Cell Ranger mkref v8.0.0 using the reference genome published by Liu et al. 2025^4^. Raw fastq reads from each caste (queen, king, soldier and worker) were aligned to the custom reference genome using Cell Ranger count v8.0.0, with default parameters. A combined cell count matrix of all the castes was then generated using Cell Ranger aggr v8.0.0, with default parameters.

Downstream analyses were performed in Python v3.12.2 using Scanpy v1.11.1^41^. Raw gene expression counts were normalized per cell using total-count normalization (sc.pp.normalize_total) to a target sum of 10,000 counts per cell and log-transformed (sc.pp.log1p). Highly variable genes were identified using sc.pp.highly_variable_genes (min_mean = 0.0125, max_mean = 3, min_disp = 0.5), and downstream dimensionality reduction was performed on this subset. Principal component analysis (PCA) was computed (sc.tl.pca, svd_solver=“arpack”) and the first 40 principal components were used to construct a k-nearest neighbour graph (k = 10). UMAP embeddings were generated, and clusters were identified using the Leiden algorithm (resolution = 0.9, random_state = 0). The embedding was coloured and visualised by cell type (**Figure 1C**), Leiden clusters (**Supplementary Figure 1B**), and caste (**Figure 2B, Supplementary Figure 1B**).

*D. melanogaster* orthologs for termite genes were identified using the BLASTX command line tool^42^ against a *D. melanogaster* protein database constructed from protein translation sequences from FlyBase (release 6.62) ^43^ (https://s3ftp.flybase.org/genomes/Drosophila_melanogaster/dmel_r6.62_FB2025_01/fasta/index.html). The top hit per query was retained with an e-value cutoff of 10^-20^.

### Cell type annotation

Automated cell type labels were obtained using SATURN-xfer^26^. Cell type annotations were cross-referenced with curated marker genes from the *D. melanogaster* single-cell atlas^22^. Eighteen clusters exhibited expression of conserved canonical markers and were therefore assigned to known *D. melanogaster* cell types. The mean expression by gene of the conserved markers were visualised using a Scanpy dotplot (**Figure 1D**). Six additional clusters (T1–T6), each defined by distinct upregulated marker genes but lacking clear correspondence to established fly cell types, were retained as novel cell types and designated T1–T6. The mean expression by gene of the marker genes were visualised using a Scanpy dotplot (**Supplementary Figure 1D**). The expression of *klu* (Znev00010248) was visualised in the stem cell (**Figure 1E**).

The Scanpy rank gene groups function (Wilcoxon rank-sum test with default parameters) was used to calculate cell type markers (logFC > 0, adjusted p-value < 0.05) for both conserved and novel cell types, as well as each leiden cluster (**Supplementary Data 1**).

Relative cell type abundance across castes was quantified. For each caste, the number of cells per cell type was counted and converted to percentages relative to the total number of cells in that caste. The percentage abundance of cell types in castes were plotted across a subset of cell types (**Figure 2C**) and across all cell types (**Supplementary Figure 2C**).

### Differential gene expression analysis

Differential expression between three castes (queen, king, and soldier) and the reference caste (worker) was calculated within each cell type using the Scanpy rank gene groups function (Wilcoxon rank-sum test with default parameters), requiring a minimum of two cells per caste, per cell type. For each cell type, significantly upregulated and downregulated genes were identified using log fold change and adjusted p-value thresholds (p-value < 0.05). The total number of upregulated and downregulated genes per cell type and caste was quantified and a subset of cell types were plotted (**Figure 2D**). The number of genes exclusively upregulated in each caste compared to workers (p-value <0.05) for each cell type was also calculated and plotted (**Figure 2E**). The total number of upregulated and downregulated, as well as caste specific upregulated genes per cell type for all cell types was also plotted (**Supplementary Figure 2D, E**).

Significant up-regulated and down-regulated gene lists for queen, king, and soldier were represented as sets and used to compute unique, pairwise-overlap, and three-way overlap categories per cell type using set operations (**Supplementary Data 2, Supplementary Data 3**). Summary counts of genes (adjusted p-value < 0.05) per overlap category were compiled per cell type and visualised using Venn diagrams for each cell type (**Supplementary Figure 3A, B**). Summary counts per overlap category of upregulated genes (adjusted p-value < 0.05) were visualised using Venn diagrams for the stem cell and muscle cell (**Figure 2F**). For each cell type and category, three significantly expressed genes belonging to each category were visualized using a Scanpy dotplot (**Figure 3E**), with gene expression values standardized across genes (standard_scale = “var”). The expression of the conserved canonical marker of fat cells, Apoltp (Znev00012632), and CG34370 (Znev00004477), a gene upregulated in all castes compared to workers, were visualised on a UMAP embedding (**Figure 3B**) and violin plot of the fat cell nuclei (**Figure 3C**). The expression of the conserved canonical marker of muscle cells, Mhc (Znev00001420), and TpnC47D (Znev00013934), a gene upregulated only in soldiers compared to workers, were visualised on a UMAP embedding (**Figure 3B**) and violin plot of the muscle cell nuclei (**Figure 3C**).

Differential expression was calculated for king, queen, and soldier relative to workers using a Wilcoxon rank-sum test as described above. For the same cell type, cell-type marker genes were identified by comparing cells of that cell type to all other cells (cell type vs rest), as above. For each gene present in both result tables, the Wilcoxon score from the caste comparison was plotted against the Wilcoxon score from the cell-type comparison. Scatter plots were generated for each selected cell type using Matplotlib. Points were coloured by the Euclidean magnitude of the combined scores (**Figure 3D, Supplementary Figure 4**).

Genes involved in juvenile hormone (JH) biosynthesis were identified based on sequences reported by Terrapon *et. al*. 2014^2^. Thirteen JH pathway genes were retrieved from the termite genome database (http://termitegenome.org/) and used as reference sequences. Homologous sequences from the *Z. nevadensis* assembly^4^ to these thirteen JH pathway genes were identified using the BLASTN command line^42^ with an e-value threshold of 1 × 10^−5^ to retain significant matches (**Supplementary Table S6**). The gene ‘gce’ (Znev00002391) was previously identified as a *D. melanogaster* ortholog (FBgn0261703). The expression of eight of these genes were visualised in the worker caste across five cell types using Scanpy (**Figure 4A)** and all castes (**Supplementary Figure 5A**). The expression level of all homologous JH pathway genes and castes were visualized across all cell types in the worker using Scanpy (**Supplementary Figure 5B-E**). Gene expression values were standardised across genes (standard_scale=‘var’), and dot size was scaled to the fraction of expressing cells with a maximum dot size of 0.2.

For each comparison between workers and another caste (queen, king, or soldier), the difference in mean expression (worker minus comparison caste) was calculated for every gene– cell type combination in which both castes were represented. For JH biosynthesis genes, these differences were pooled across all genes in the pathway, whereas *gce* was analysed separately across cell types. Statistical significance was assessed using a two-sided Wilcoxon signed-rank test testing whether the median expression difference deviated from zero (**Supplementary Table S7, S8**).

To visualise the distribution of expression differences, empirical cumulative distribution functions (ECDFs) (**Supplementary Figure 5F, G**) and kernel density estimates (KDEs) (**Supplementary Figure 5F, G**) were generated for each caste comparison.

Vitellogenin (Vg) genes were downloaded from the NCBI *Z. nevadensis* genome ZooNev1.0 (GCF_000696155.1). Homologous sequences from the *Z. nevadensis* assembly^4^ were identified using the BLASTN command line^42^ with an e-value threshold of 1 × 10^−5^ to retain significant matches (**Supplementary Table S9**). The expression level of Vg genes was visualized across all cell types in all castes using Scanpy (**Supplementary Figure 6A-D**). The expression of eight of these genes was visualised across all castes and across three cell types using Scanpy (**Figure 4B)**. Dot size was scaled to the fraction of expressing cells with a maximum dot size of 0.2.

The cell-type abundance and gene expression across termite castes of ‘Vg T1’ (Znev00005980) in fat cells were quantified (**Figure 4C**). The percentage abundance of the fat cells in each caste was computed by dividing the number of cells of that type by the total number of cells for each caste and multiplying by 100. The mean log-normalised expression values (as calculated above) were averaged across fat cells for each caste.

Genes involved in immunity were identified based on sequences reported by Terrapon et al. 2014^2^. One-hundred and seventy immune system genes were downloaded from the NCBI *Z. nevadensis* genome ZooNev1.0 (GCF_000696155.1). Homologous sequences from the *Z. nevadensis* assembly^4^ were identified using the BLASTN command line tool^42^ with an e-value threshold of 1 × 10^−5^ to retain significant matches. A subset of six transcripts (PGRP, GNBP, cSP, Easter, Toll-like receptor, and spaetzle) (**Supplementary Table S10**) were then visualised in the following ways. To quantify the number of different cell types that were expressing these transcripts, a cell type was counted for a given gene and caste if at least 10% of cells in that cell type were expressing the gene with a log mean expression of at least 0.01. Results were visualised as a grouped bar chart (**Figure 4D**). The expression of PGRP and the Toll-like receptor was visualised across all nuclei on a UMAP embedding (**Figure 4E**). The percentage abundance of the crystal cells in each caste was computed by dividing the number of cells of that type by the total number of cells for each caste and multiplying by 100 (**Figure 4G**). The expression level of these transcripts was visualized on a UMAP embedding of crystal cells and hemocytes (**Figure 4H**). The expression level of these transcripts was visualized on a dot plot of crystal cells across castes (**Figure 4H**), with colour intensity capped at an expression value of 1.5 (vmax=1.5) and dot sizes scaled to the fraction of expressing cells with a maximum dot size of 0.4. The expression level of these transcripts was also visualized on a dot plot of across castes and all cell types (**Supplementary Figure 8A-D**).

Carbohydrate-Active Enzyme (CAZymes) transcripts were identified in 45 termite species and two cockroaches^4^. To visualise CAZyme expression, mean expression was calculated per transcript across cell types. The expression of the top 3 transcripts (ranked by maximum mean expression across cell types) were visualised across castes (queen, king, soldier, and worker) and four cell types using dotplots (**Figure 4I**), with colour intensity capped at an expression value of 4.5 (vmax=4.5). The expression level of these transcripts was also visualized on a dot plot across castes and cell types (**Supplementary Figure 9A-D**). The top 20 genes (ranked by maximum mean expression across cell types) were selected and expression was visualised across castes (**Supplementary Figure 9E**).

Hierarchical orthogroups (HOGs) were identified in 45 termite species and two cockroaches^4^. For each HOG, multiple sequence alignment of proteins were generated using MAFFT v7.508^44^ with the –auot option, and converted into codon alignment using PAL2NAL v14 ^45^. From the codon alignment, pairwise synonymous and nonsynonymous substitution rate (Ks and Ka) was computed using the *kaks* function with *rmgap*=FALSE implemented in R pakage *seqinr*^*46*^, using the methods of Li (1993) and the substitution model of Kimura (1980). From these pairwise Ka and Ks values, we extracted the comparison of *Hodotermopsis* and *Zootermopsis*, and only kept HOGs that were single-copy in both species.

Genes with a Ka or Ks value greater than three were removed from subsequent analysis. The distribution of Ka/Ks ratios for genes represented in the single-cell dataset was visualized using a histogram (**Supplementary Figure S10A**). The relationship between Ka and Ks was visualized using a scatter plot, with a diagonal reference line representing Ka = Ks (**Supplementary Figure S10B**). Genes were grouped into four categories: missing dN/dS values, dN/dS < 0.05, 0.05–1.1, and dN/dS > 1.1. The proportion of all genes in the dataset belonging to each category was visualised as a bar plot (**Supplementary Figure S10C**).

For each gene, the number of cell types in which it was expressed was calculated. A gene was considered expressed in a cell type if mean log-normalised expression was ≥ 0.1 and at least 10% of cells of that type had detectable expression (expression > 0). The distribution of the number of expressing cell types in each dN/dS category was visualised using complementary CDF (**Supplementary Figure S10D**) and KDE (**Supplementary Figure S10E**) plots. The distribution of the number of expressing cell types in each dN/dS category was visualised using complementary CDF without the genes missing dN/dS values (**Figure 4J**).

## Supplementary Information

### Supplementary Information

Supplementary Tables 1-10, Supplementary Figures 1-10

### Supplementary Data 1

Cell-type markers.

### Supplementary Data 2

Significant up-regulated gene lists (adjusted p-value < 0.05) for queen, king, and soldier castes.

### Supplementary Data 3

Significant down-regulated gene lists (adjusted p-value < 0.05) for queen, king, and soldier castes.

