## Supplementary Information for "Social caste switching triggers emergence of novel cellular identities in *Zootermopsis* termites"

#### Supplementary Tables

**Table S1: Total cells sequenced per caste.**

| | Queen | King | Soldier | Worker |
|---|---|---|---|---|
| **Total number of cells** | 6529 | 2288 | 4805 | 10630 |

**Table S2: Relative cell type abundance across castes across all cells.** Number of cells in each caste and cell type, as well as the relative abundance of that cell type within each caste. P-values were calculated using Fisher's exact test, where 'reproductive castes' are defined as queen and king, and non-reproductive castes are soldier and worker. Significance is defined as $p < 0.05$ and coloured red. P-values were reported to two decimal places.

| | Queen | | King | | Soldier | | Worker | | Reproductive (Q+K) | Non-reproductive (W+S) | Enriched in |
|---|---|---|---|---|---|---|---|---|---|---|---|
| **Cell Type** | **n** | **%** | **n** | **%** | **n** | **%** | **n** | **%** | | | |
| fat cell | 1105 | 16.9 | 226 | 9.9 | 344 | 7.2 | 543 | 5.1 | 0 | 1 | reproductive |
| stem cell | 1144 | 17.5 | 287 | 12.5 | 387 | 8.1 | 838 | 7.9 | 0 | 1 | reproductive |
| intestinal progenitor like | 895 | 13.7 | 231 | 10.1 | 320 | 6.7 | 784 | 7.4 | 0 | 1 | reproductive |
| hemocyte | 323 | 4.9 | 62 | 2.7 | 212 | 4.4 | 187 | 1.8 | 0 | 1 | reproductive |
| crystal cell | 192 | 2.9 | 32 | 1.4 | 199 | 4.1 | 129 | 1.2 | 0.02 | 0.98 | reproductive |
| cardiac cell | 275 | 4.2 | 81 | 3.5 | 77 | 1.6 | 89 | 0.8 | 0 | 1 | reproductive |
| antimicrobial | 113 | 1.7 | 29 | 1.3 | 17 | 0.4 | 28 | 0.3 | 0 | 1 | reproductive |
| CNS neuron | 68 | 1 | 28 | 1.2 | 27 | 0.6 | 61 | 0.6 | 0 | 1 | reproductive |
| T1 | 0 | 0 | 160 | 7 | 1 | 0 | 0 | 0 | 0 | 1 | reproductive |
| female | 271 | 4.2 | 0 | 0 | 10 | 0.2 | 14 | 0.1 | 0 | 1 | reproductive |

| | | | | | | | | | | | |
|---|---|---|---|---|---|---|---|---|---|---|---|
| reproductive system | | | | | | | | | | | |
| epithelial | 496 | 7.6 | 337 | 14.7 | 669 | 13.9 | 1544 | 14.5 | 1 | 0 | non-reproductive |
| muscle cell | 179 | 2.7 | 168 | 7.3 | 734 | 15.3 | 1823 | 17.1 | 1 | 0 | non-reproductive |
| T3 | 228 | 3.5 | 139 | 6.1 | 291 | 6.1 | 576 | 5.4 | 1 | 0 | non-reproductive |
| neuron | 168 | 2.6 | 115 | 5 | 378 | 7.9 | 1288 | 12.1 | 1 | 0 | non-reproductive |
| T6 | 137 | 2.1 | 80 | 3.5 | 129 | 2.7 | 494 | 4.6 | 1 | 0 | non-reproductive |
| sensory neuron | 103 | 1.6 | 57 | 2.5 | 224 | 4.7 | 440 | 4.1 | 1 | 0 | non-reproductive |
| glial cell | 136 | 2.1 | 56 | 2.4 | 258 | 5.4 | 321 | 3 | 1 | 0 | non-reproductive |
| T4 | 90 | 1.4 | 44 | 1.9 | 110 | 2.3 | 267 | 2.5 | 1 | 0 | non-reproductive |
| oenocyte | 23 | 0.4 | 8 | 0.3 | 100 | 2.1 | 142 | 1.3 | 1 | 0 | non-reproductive |
| excretory | 3 | 0 | 1 | 0 | 59 | 1.2 | 115 | 1.1 | 1 | 0 | non-reproductive |
| Malpighian tubule stellate | 13 | 0.2 | 1 | 0 | 4 | 0.1 | 38 | 0.4 | 0.97 | 0.05 | non-reproductive |
| T2 | 220 | 3.4 | 73 | 3.2 | 58 | 1.2 | 465 | 4.4 | 0.62 | 0.41 | ns |
| T5 | 219 | 3.4 | 57 | 2.5 | 141 | 2.9 | 289 | 2.7 | 0.07 | 0.94 | ns |
| gustatory neuron | 128 | 2 | 16 | 0.7 | 56 | 1.2 | 155 | 1.5 | 0.06 | 0.96 | ns |

**Table S3: Upregulated genes in castes per cell type.** Differential expression analysis was run using Wilcoxon DE for each cell type and caste. Genes with a LogFC > 0 and an adjusted p-value < 0.05 were counted for all categories using set overlaps. S = all genes significant in soldier vs worker. K = all genes significant in king vs worker. Q = all genes significant in queen vs worker. S\(K∪Q) = only significant in soldier, not in king or queen. K\(S∪Q) = only significant in king, not in soldier or queen. Q\(S∪K) = only significant in queen, not in soldier or king. (S∩K)\Q = significant in soldier and king, but not queen. (S∩Q)\K = significant in soldier and queen, but not king. (K∩Q)\S = significant in king and queen, but not soldier. S∩K∩Q = significant in all three castes. S∪K∪Q = significant in at least one caste.

| **N. of Up-Reg Genes** | **S** | **K** | **Q** | **S\(K∪Q) Unique Soldier** | **K\(S∪Q) Unique King** | **Q\(S∪K) Unique Queen** | **(S∩K)\Q Soldier and King only** | **(S∩Q)\K Soldier and Queen only** | **(K∩Q)\S Kig and Queen only** | **S∩K∩Q All Q, K and S** | **S∪K∪Q Any Q, K, S** |
|---|---|---|---|---|---|---|---|---|---|---|---|
| **Cell Type** | | | | | | | | | | | |
| fat cell | 726 | 236 | 531 | 349 | 39 | 166 | 46 | 214 | 34 | 117 | 965 |
| intestinal progenitor like | 407 | 337 | 1098 | 64 | 46 | 669 | 23 | 161 | 109 | 159 | 1231 |
| epithelial | 399 | 166 | 246 | 218 | 17 | 88 | 42 | 51 | 19 | 88 | 523 |
| muscle cell | 378 | 46 | 43 | 330 | 12 | 15 | 20 | 14 | 0 | 14 | 405 |
| oenocyte | 185 | 0 | 10 | 178 | 0 | 3 | 0 | 7 | 0 | 0 | 188 |
| neuron | 175 | 18 | 25 | 157 | 6 | 11 | 4 | 6 | 0 | 8 | 192 |
| stem cell | 124 | 144 | 509 | 27 | 27 | 354 | 3 | 41 | 61 | 53 | 566 |
| glial cell | 48 | 11 | 26 | 37 | 4 | 18 | 4 | 5 | 1 | 2 | 71 |
| sensory neuron | 39 | 5 | 35 | 29 | 1 | 23 | 0 | 8 | 2 | 2 | 65 |

| | | | | | | | | | | | |
|---|---|---|---|---|---|---|---|---|---|---|---|
| T3 | 26 | 25 | 285 | 17 | 10 | 266 | 1 | 5 | 11 | 3 | 313 |
| T6 | 26 | 4 | 32 | 19 | 2 | 24 | 0 | 6 | 1 | 1 | 53 |
| gustatory neuron | 22 | 0 | 32 | 17 | 0 | 27 | 0 | 5 | 0 | 0 | 49 |
| excretory | 22 | 0 | 0 | 22 | 0 | 0 | 0 | 0 | 0 | 0 | 22 |
| T5 | 20 | 7 | 20 | 17 | 4 | 16 | 0 | 1 | 1 | 2 | 41 |
| hemocyte | 20 | 6 | 37 | 7 | 0 | 27 | 3 | 7 | 0 | 3 | 47 |
| crystal cell | 19 | 0 | 44 | 13 | 0 | 38 | 0 | 6 | 0 | 0 | 57 |
| cardiac cell | 13 | 4 | 27 | 9 | 2 | 22 | 0 | 3 | 1 | 1 | 38 |
| T2 | 10 | 14 | 62 | 5 | 5 | 51 | 0 | 2 | 6 | 3 | 72 |
| T4 | 4 | 1 | 10 | 3 | 0 | 9 | 0 | 0 | 0 | 1 | 13 |
| T1 | 0 | 0 | 0 | 0 | 0 | 0 | 0 | 0 | 0 | 0 | 0 |
| antimicrobial | 0 | 0 | 13 | 0 | 0 | 13 | 0 | 0 | 0 | 0 | 13 |
| CNS neuron | 0 | 0 | 4 | 0 | 0 | 4 | 0 | 0 | 0 | 0 | 4 |
| Malpighian tubule stellate | 0 | 0 | 0 | 0 | 0 | 0 | 0 | 0 | 0 | 0 | 0 |
| female reproductive system | 0 | 0 | 0 | 0 | 0 | 0 | 0 | 0 | 0 | 0 | 0 |

**Table S4: Downregulated genes in castes per cell type.** Differential expression analysis was run using Wilcoxon DE for each cell type and caste. Genes with a LogFC < 0 and an adjusted p-value < 0.05 were counted for all categories using set overlaps. S = all genes significant in soldier vs worker. K = all genes significant in king vs worker. Q = all genes significant in queen vs worker. S\(K∪Q) = only significant in soldier, not in king or queen. K\(S∪Q) = only significant in king, not in soldier or queen. Q\(S∪K) = only significant in queen, not in soldier or king. (S∩K)\Q = significant in soldier and king, but not queen. (S∩Q)\K = significant in soldier and queen, but not king. (K∩Q)\S = significant in king and queen, but not soldier. S∩K∩Q = significant in all three castes. S∪K∪Q = significant in at least one caste.

| **N. of Down-Reg Genes** | S | K | Q | S\(K∪Q) Unique Soldier | K\(S∪Q) Unique King | Q\(S∪K) Unique Queen | (S∩K)\Q Soldier and King only | (S∩Q)\K Soldier and Queen only | (K∩Q)\S Kig and Queen only | S∩K∩Q All Q, K and S | S∪K∪Q Any Q, K, S |
|---|---|---|---|---|---|---|---|---|---|---|---|
| **Cell Type** | | | | | | | | | | | |
| fat cell | 194 | 162 | 284 | 47 | 24 | 101 | 10 | 55 | 46 | 82 | 365 |
| intestinal progenitor like | 176 | 65 | 112 | 93 | 12 | 37 | 17 | 39 | 9 | 27 | 234 |
| muscle cell | 97 | 8 | 12 | 87 | 2 | 5 | 3 | 4 | 0 | 3 | 104 |
| stem cell | 68 | 20 | 44 | 43 | 3 | 16 | 2 | 13 | 5 | 10 | 92 |
| epithelial | 45 | 28 | 31 | 21 | 8 | 5 | 1 | 7 | 3 | 16 | 61 |
| neuron | 41 | 6 | 18 | 29 | 0 | 5 | 0 | 7 | 1 | 5 | 47 |
| oenocyte | 26 | 0 | 3 | 24 | 0 | 1 | 0 | 2 | 0 | 0 | 27 |
| sensory neuron | 21 | 2 | 6 | 15 | 0 | 0 | 0 | 4 | 0 | 2 | 21 |

| | | | | | | | | | | | |
|---|---|---|---|---|---|---|---|---|---|---|---|
| glial cell | 16 | 1 | 4 | 14 | 0 | 2 | 0 | 1 | 0 | 1 | 18 |
| T6 | 15 | 1 | 8 | 10 | 0 | 3 | 0 | 4 | 0 | 1 | 18 |
| T3 | 12 | 8 | 21 | 5 | 2 | 12 | 0 | 3 | 2 | 4 | 28 |
| gustatory neuron | 10 | 0 | 16 | 4 | 0 | 10 | 0 | 6 | 0 | 0 | 20 |
| T5 | 9 | 5 | 13 | 4 | 1 | 8 | 1 | 2 | 1 | 2 | 19 |
| T2 | 9 | 3 | 18 | 5 | 0 | 14 | 0 | 1 | 0 | 3 | 23 |
| excretory | 9 | 0 | 0 | 9 | 0 | 0 | 0 | 0 | 0 | 0 | 9 |
| T4 | 7 | 0 | 2 | 5 | 0 | 0 | 0 | 2 | 0 | 0 | 7 |
| hemocyte | 6 | 1 | 7 | 2 | 0 | 3 | 0 | 3 | 0 | 1 | 9 |
| cardiac cell | 6 | 1 | 8 | 3 | 0 | 4 | 0 | 3 | 1 | 0 | 11 |
| crystal cell | 4 | 0 | 8 | 1 | 0 | 5 | 0 | 3 | 0 | 0 | 9 |
| T1 | 0 | 0 | 0 | 0 | 0 | 0 | 0 | 0 | 0 | 0 | 0 |
| antimicrobial | 0 | 0 | 6 | 0 | 0 | 6 | 0 | 0 | 0 | 0 | 6 |
| CNS neuron | 0 | 0 | 2 | 0 | 0 | 2 | 0 | 0 | 0 | 0 | 2 |
| Malpighian tubule stellate | 0 | 0 | 0 | 0 | 0 | 0 | 0 | 0 | 0 | 0 | 0 |
| female reproductive system | 0 | 0 | 0 | 0 | 0 | 0 | 0 | 0 | 0 | 0 | 0 |

**Table S5: Cell markers of cell type T1.**

This table lists the 10 highest ranked upregulated genes for cell type T1, ranked by Wilcoxon score. Differential expression was performed using a Wilcoxon rank-sum test to identify cell-type-specific markers across all cells, independent of caste. Genes with logFC > 0 and adjusted p-value < 0.05 were considered significant.

| Gene | Score | P-value | logFC | *D. melanogaster* ortholog |
|---|---|---|---|---|
| Znev00006597 | 21.60052681 | 1.7758E-103 | 11.44482803 | |
| Znev00002920 | 21.310009 | 9.1679E-101 | 10.55583668 | FASN3 |
| Znev00003603 | 21.18513107 | 1.3096E-99 | 10.63030338 | |
| Znev00012628 | 20.98117447 | 9.74568E-98 | 8.730919838 | L |
| Znev00006383 | 20.8059845 | 3.82031E-96 | 11.80536461 | |
| Znev00008233 | 20.6657238 | 7.04916E-95 | 11.8903265 | SPE |
| Znev00005648 | 20.35894775 | 3.86797E-92 | 6.18835783 | apolpp |
| Znev00010080 | 19.82352638 | 1.86566E-87 | 7.657146931 | CCAP-R |
| Znev00013765 | 19.4005928 | 7.6288E-84 | 3.789125919 | Srrm234 |
| Znev00004888 | 17.97098923 | 3.28802E-72 | 5.148839951 | |

**Table S6: JH biosynthesis pathway genes**

| Znev ID | JH Gene |
|---|---|
| Znev00000780 | 3-hydroxy-3-methylglutaryl-CoA reductase (HMGR) |
| Znev00004071 | 3-hydroxy-3-methylglutaryl-CoA synthase 2 (HMGS2) |
| Znev00004155 | isopentenyl diphosphate isomerase (IPPI) |
| Znev00004225 | JH epoxidase (CYP15A1) |
| Znev00004645 | JH epoxidase (CYP15A1) |
| Znev00004952 | JH acid methyltransferase (JHAMT) |
| Znev00005476 | 3-hydroxy-3-methylglutaryl-CoA synthase 1 (HMGS1) |
| Znev00006447 | mevalonate kinase (MK) |
| Znev00010869 | farnesal dehydrogenase 1 (FD1) |
| Znev00011682 | phosphomevalonate kinase (PK) |
| Znev00011953 | acetoacetyl-CoA thiolase (AcoAT) |
| Znev00013778 | farnesal dehydrogenase 2 (FD2) |
| Znev00013967 | farnesyl pyrophosphate synthase (FPS) |

| | |
|---|---|
| Znev00014035 | diphosphomevalonate decarboxylase (DD) |
| Znev00002391 | Germ cell-expressed bHLH-PAS, isoform C (gce) |

**Table S7: Expression of JH biosynthesis pathway (excluding Znev00002391, gce).**

| Comparison | No. Genes x Cell Types | Median worker minus other caste | P value |
|---|---|---|---|
| Worker vs queen | 322 | -0.0128 | 0.0 |
| Worker vs king | 308 | 0 | 0.0 |
| Worker vs soldier | 322 | 0 | 0.0 |

**Table S8: Expression of Znev00002391, gce.**

| Comparison | No. Genes x Cell Types | Median worker minus other caste | P value |
|---|---|---|---|
| Worker vs queen | 23 | -0.0168 | 0.85 |
| Worker vs king | 22 | -0.0023 | 0.52 |
| Worker vs soldier | 23 | 0.0922 | 0.02 |

**Table S9: Vitellogenin (Vg) genes**

| Znev_ID | Gene_name |
|---|---|
| Znev00001161 | yl |
| Znev00005980 | Vg T1 |
| Znev00005602 | Vg T2 |
| Znev00013231 | Vg-1 T1 |
| Znev00004585 | Vg-1 T2 |
| Znev00013232 | Vg-1 T3 |

**Table S10: Toll sIgnalling pathway genes**

| Znev_ID | Gene |
|---|---|
| Znev00002332 | PGRP |
| Znev00007353 | GNBP |
| Znev00011578 | cSP (stubble) |
| Znev00008234 | Easter |
| Znev00006305 | Toll-like receptor |

| | |
|---|---|
| Znev00006857 | spaetzle |

### Supplementary Figures

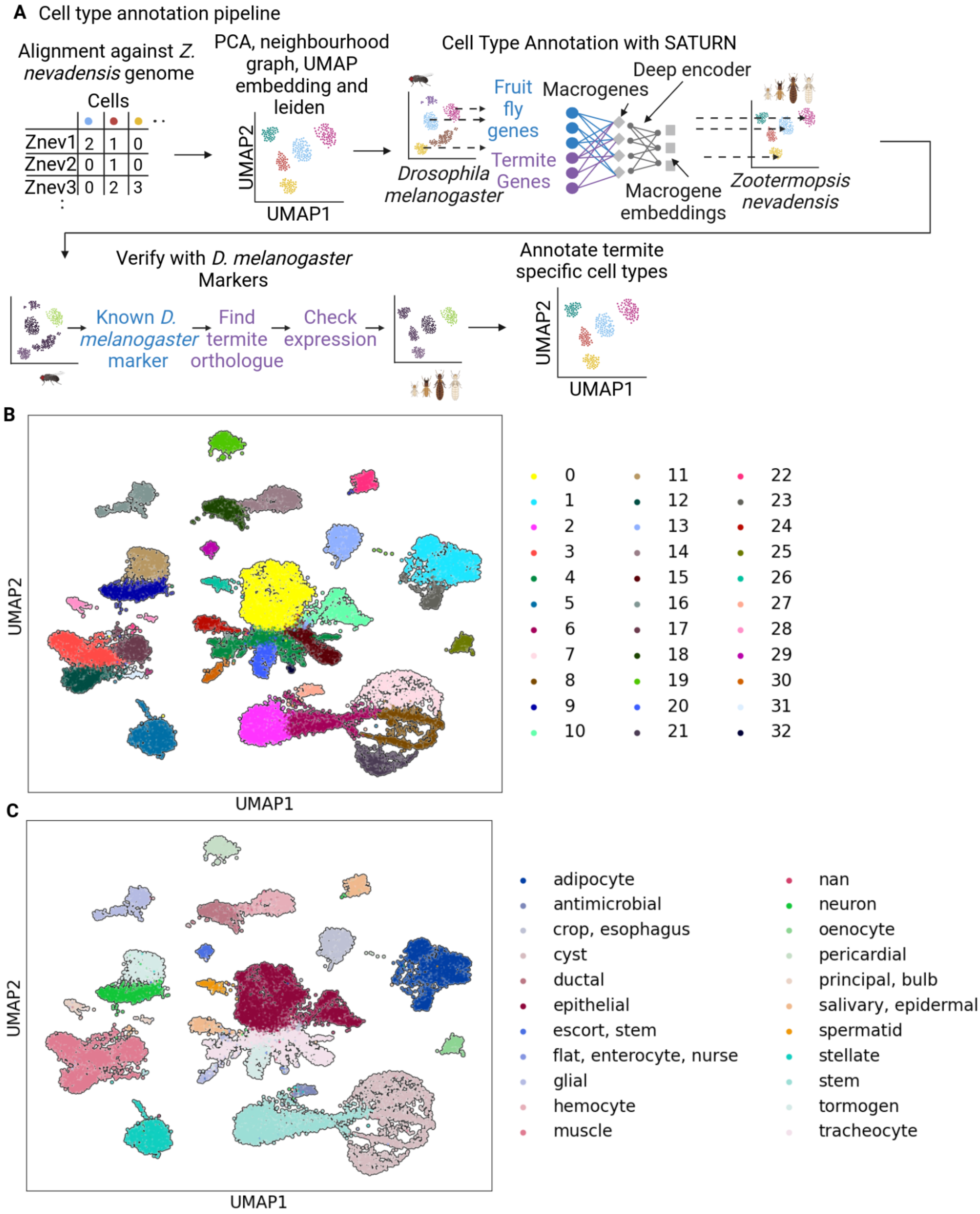

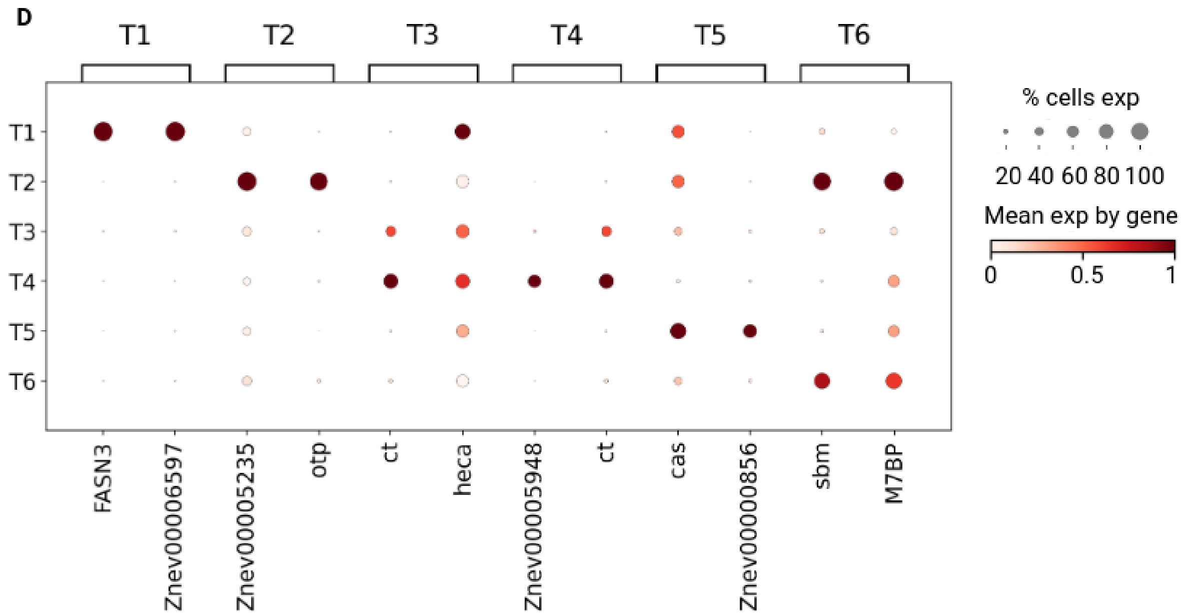


**Figure S1: Cell type annotation of cell atlas of the eusocial termite *Zootermopsis nevadensis*. A)** Overview of the cell type annotation workflow. **B)** UMAP embedding of all nuclei coloured by cell type as assigned by leiden clustering. **C)** UMAP embedding of all nuclei coloured by cell type as assigned by SATURN. **D)** Dot plot showing the expression of cell type markers across novel termite cell types. Dot size represents the proportion of cells expressing the gene and colour (white to dark red) shows average expression in that cell type.

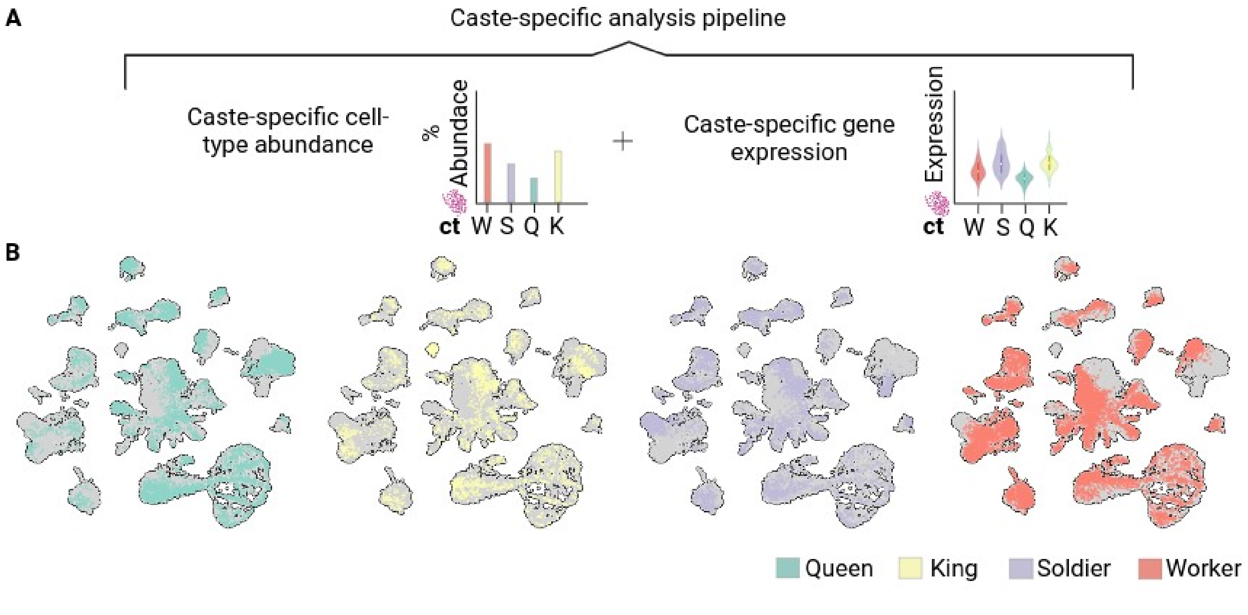

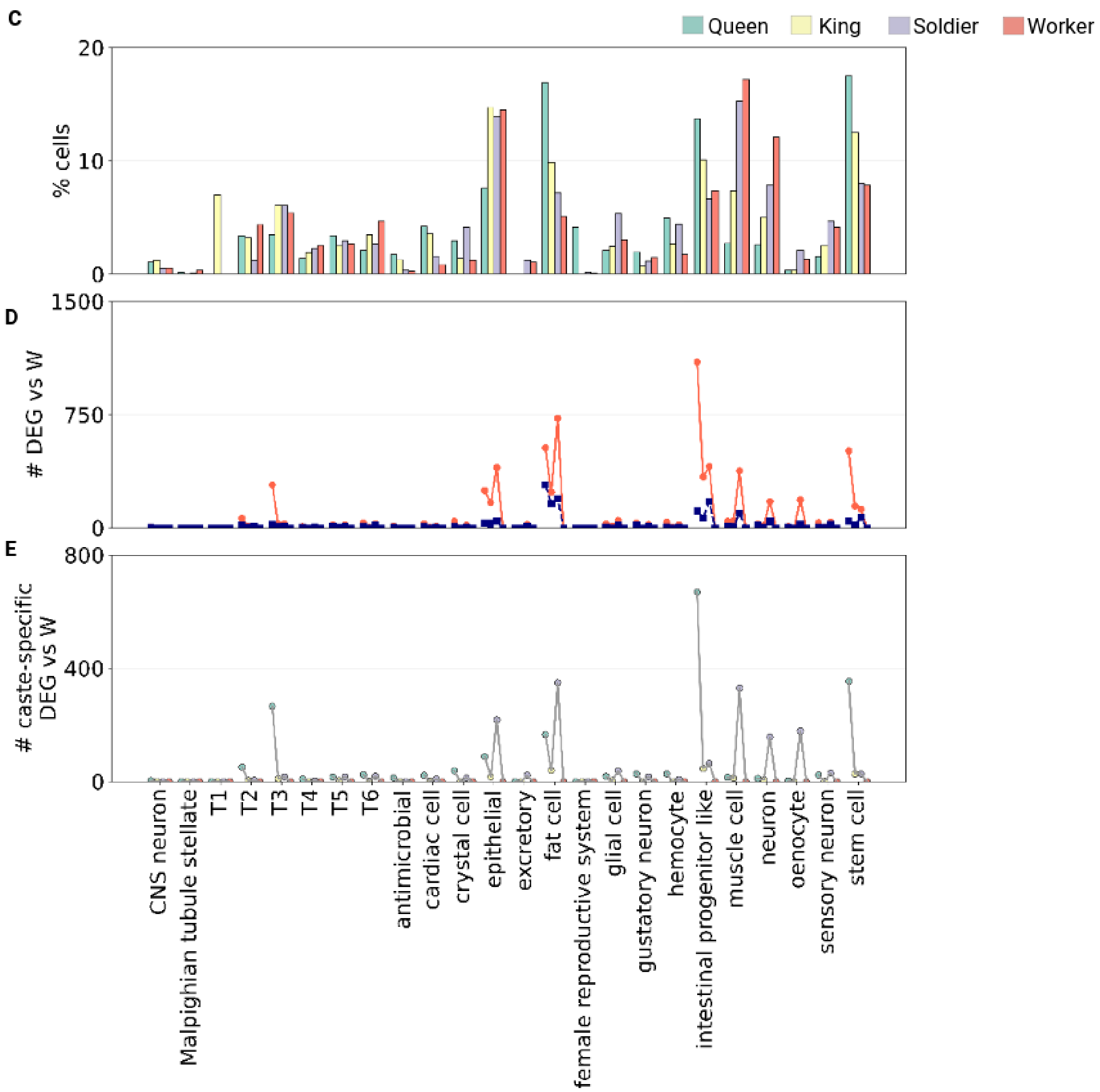


**Figure S2: Caste development analysis. A)** Diagram of the different analyses comparing castes at the single-cell level. **B)** UMAP embedding of termite nuclei as in Figure 1C, S1B, S1C, coloured by caste of origin. **C)** Relative cell type abundance across castes across all cells. **D)** Number of upregulated and downregulated genes in each caste compared to workers (adjusted p-value < 0.05), for each cell type. **E)** The number of genes exclusively upregulated in each caste compared to workers (p-value <0.05) for each cell type.

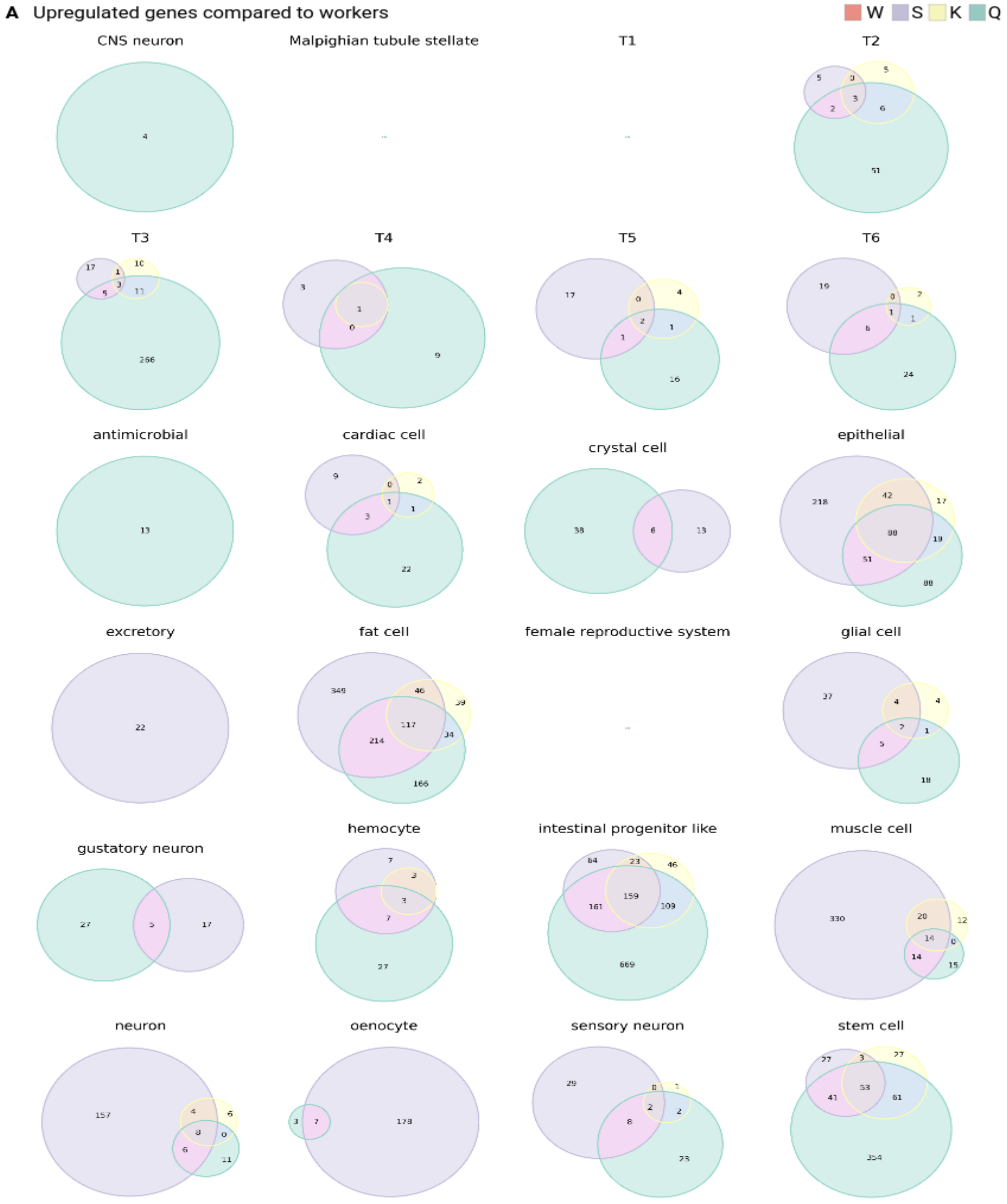
A Upregulated genes compared to workers
W S K Q
CNS neuron
Malpighian tubule stellate
T1
T2
T3
T4
T5
T6
antimicrobial
cardiac cell
crystal cell
epithelial
excretory
fat cell
female reproductive system
glial cell
gustatory neuron
hemocyte
intestinal progenitor like
muscle cell
neuron
oenocyte
sensory neuron
stem cell

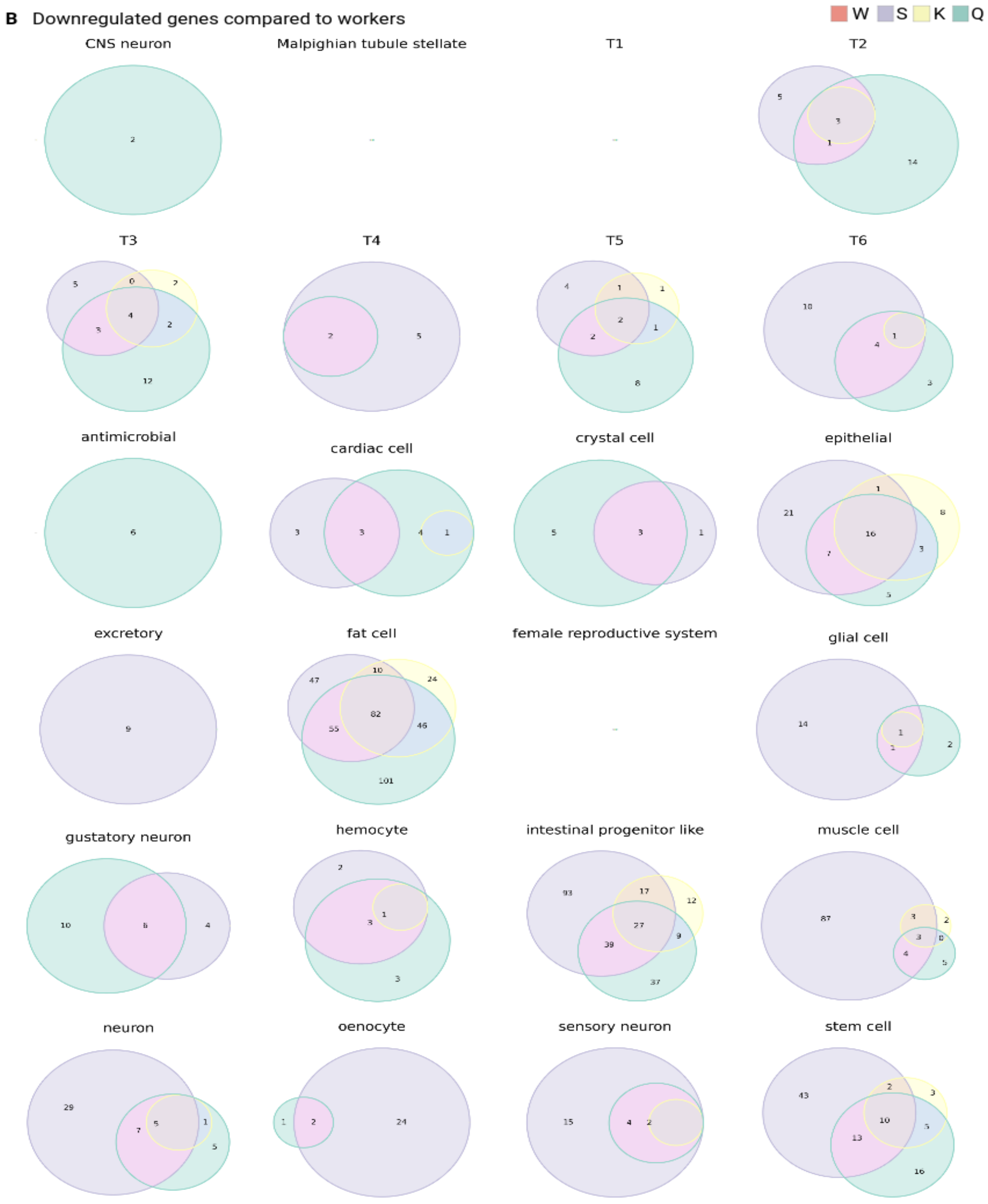
B Downregulated genes compared to workers
W S K Q
CNS neuron
Malpighian tubule stellate
T1
T2
T3
T4
T5
T6
antimicrobial
cardiac cell
crystal cell
epithelial
excretory
fat cell
female reproductive system
glial cell
gustatory neuron
hemocyte
intestinal progenitor like
muscle cell
neuron
oenocyte
sensory neuron
stem cell

**Figure S3: Differentially expressed genes across different castes and cell types. A)** Venn diagram of upregulated genes against workers in queen, king, and soldier in all cell types. **B)** Venn diagram of downregulated genes against workers in queen, king, and soldier in all cell types.

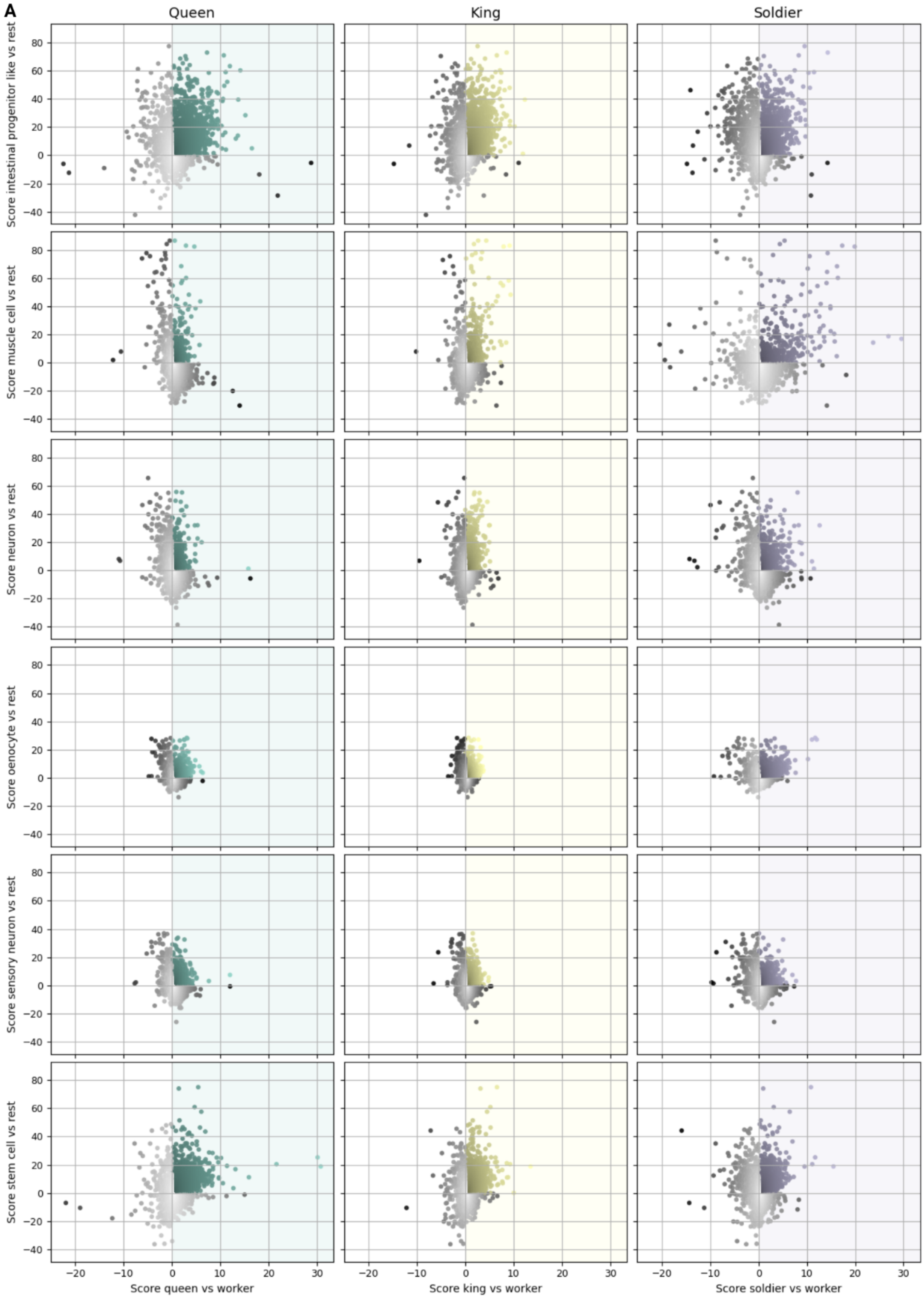

A
Queen
King
Soldier
Score intestinal progenitor like vs rest
Score muscle cell vs rest
Score neuron vs rest
Score oenocyte vs rest
Score sensory neuron vs rest
Score stem cell vs rest
Score queen vs worker
Score king vs worker
Score soldier vs worker

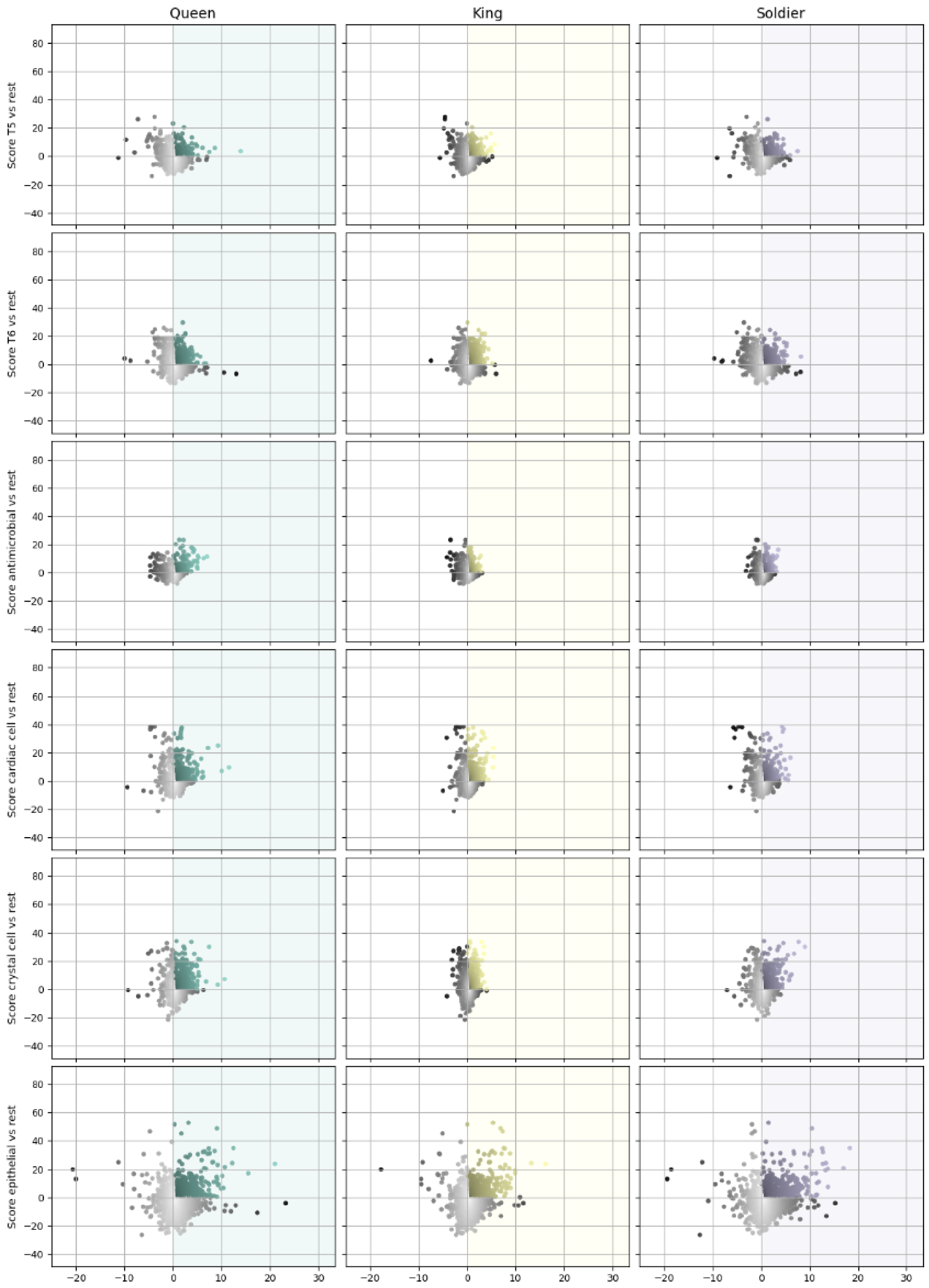

Queen
King
Soldier
Score T5 vs rest
Score T6 vs rest
Score antimicrobial vs rest
Score cardiac cell vs rest
Score crystal cell vs rest
Score epithelial vs rest
80
60
40
20
0
−20
−40
−20
−10
0
10
20
30

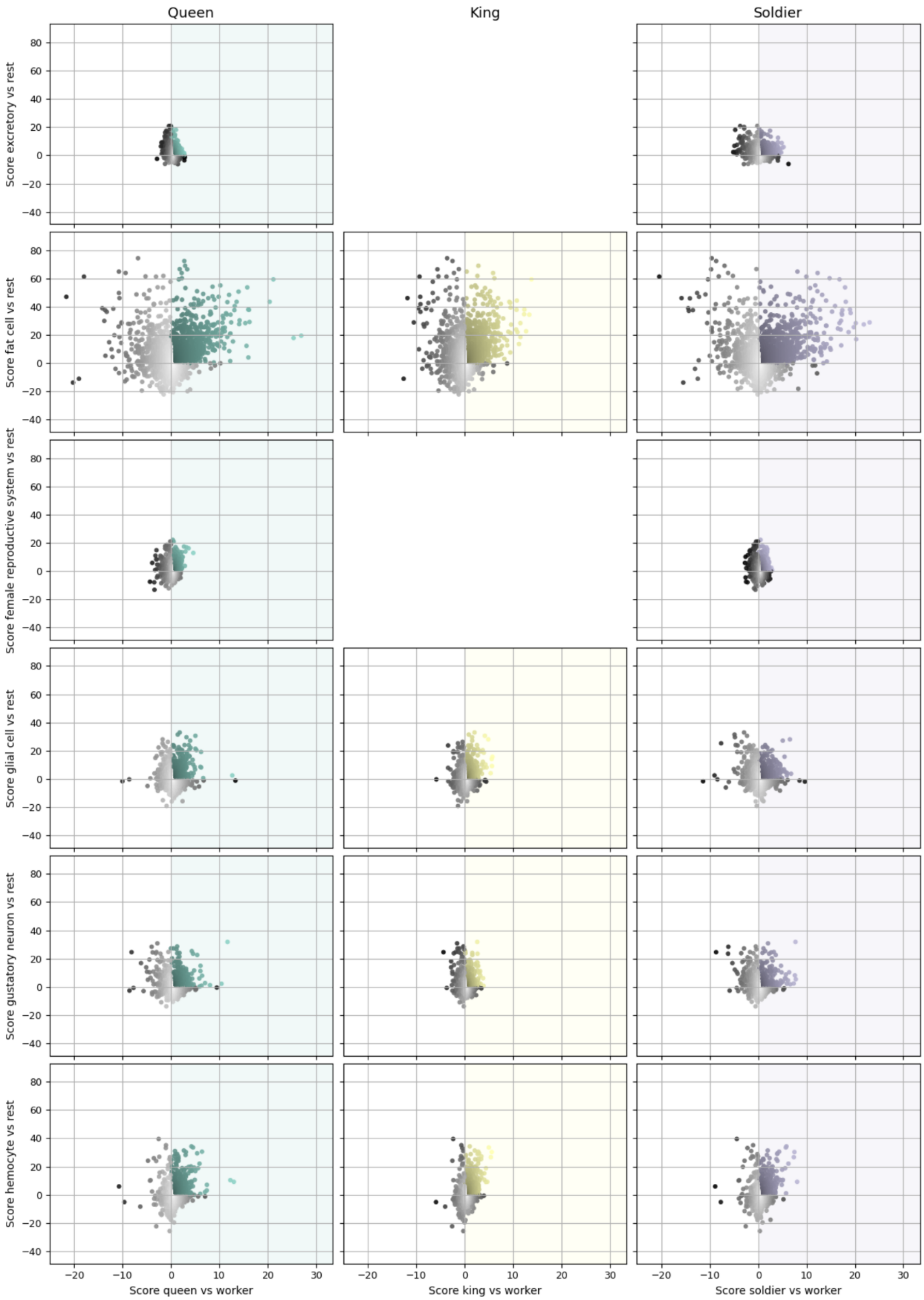

Queen
King
Soldier
Score excretory vs rest
Score fat cell vs rest
Score female reproductive system vs rest
Score glial cell vs rest
Score gustatory neuron vs rest
Score hemocyte vs rest
Score queen vs worker
Score king vs worker
Score soldier vs worker

**Figure S4: Cell-type and caste specific regulation. A)** Scatter plots for dual differential expression analysis. For each gene, represented by a dot, the x coordinate shows the statistical test score of each caste (queen, king, soldier) against worker cells of the same type. The y coordinate shows the statistical score of cells of that type versus other cells across all samples. Genes in the top right section are dual specialised.

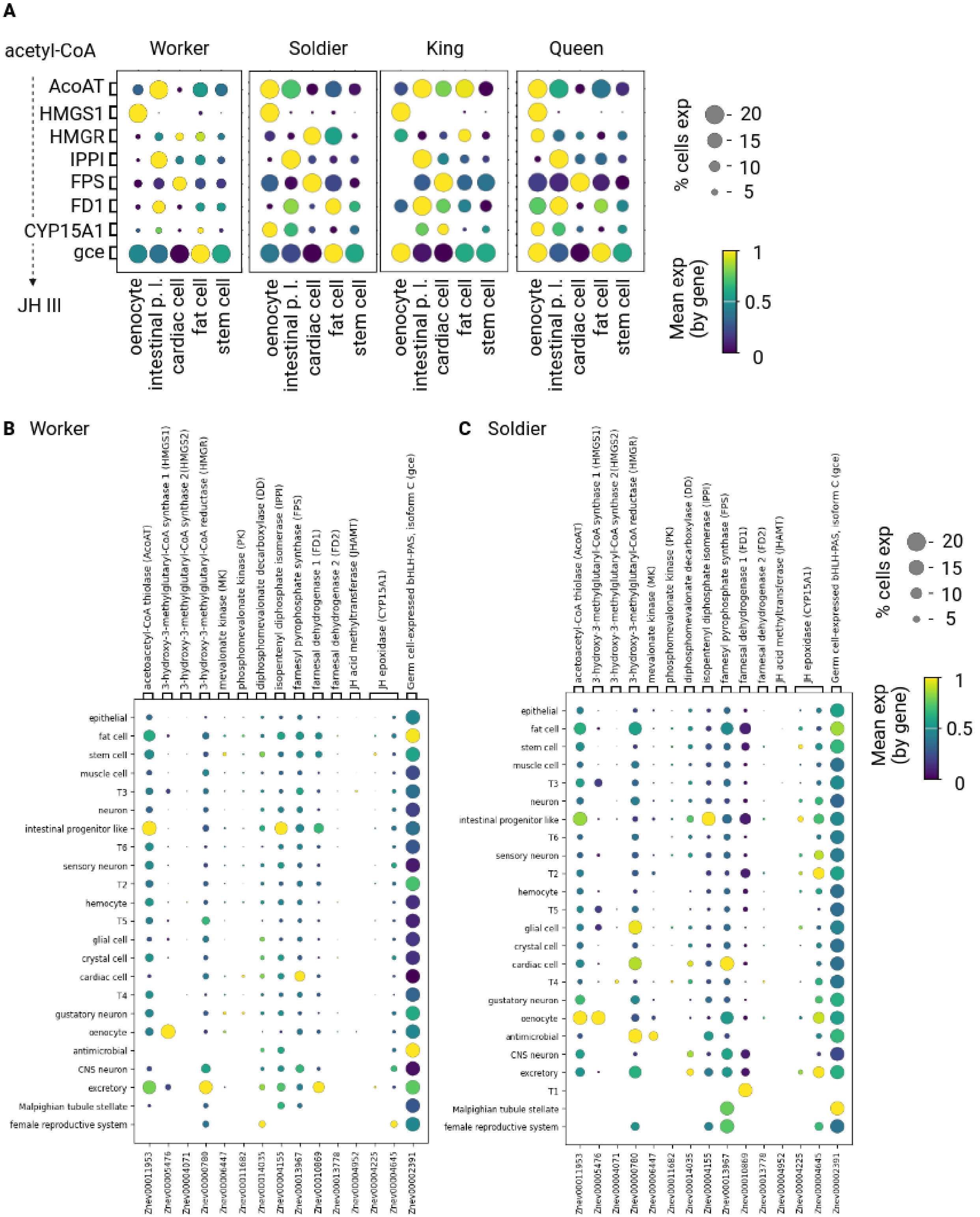

A
acetyl-CoA
JH III
Worker
Soldier
King
Queen
AcoAT
HMGS1
HMGR
IPPI
FPS
FD1
CYP15A1
gce
oenocyte
intestinal p. l.
cardiac cell
fat cell
stem cell
% cells exp
20
15
10
5
Mean exp (by gene)
1
0.5
0
B Worker
C Soldier
acetoacetyl-CoA thiolase (AcoAT)
3-hydroxy-3-methylglutaryl-CoA synthase 1 (HMGS1)
3-hydroxy-3-methylglutaryl-CoA synthase 2(HMGS2)
3-hydroxy-3-methylglutaryl-CoA reductase (HMGR)
mevalonate kinase (MK)
phosphomevalonate kinase (PK)
diphosphomevalonate decarboxylase (DD)
isopentenyl diphosphate isomerase (IPPI)
farnesyl pyrophosphate synthase (FPS)
farnesal dehydrogenase 1 (FD1)
farnesal dehydrogenase 2 (FD2)
JH acid methyltransferase (JHAMT)
JH epoxidase (CYP15A1)
Germ cell-expressed bHLH-PAS, isoform C (gce)
epithelial
fat cell
stem cell
muscle cell
T3
neuron
intestinal progenitor like
T6
sensory neuron
T2
hemocyte
T5
glial cell
crystal cell
cardiac cell
T4
gustatory neuron
oenocyte
antimicrobial
CNS neuron
excretory
T1
Malpighian tubule stellate
female reproductive system
Znev00011953
Znev00005476
Znev00004071
Znev00000780
Znev00006447
Znev00011682
Znev00014035
Znev00004155
Znev00013967
Znev00010869
Znev00013778
Znev00004952
Znev00004225
Znev00004645
Znev00002391

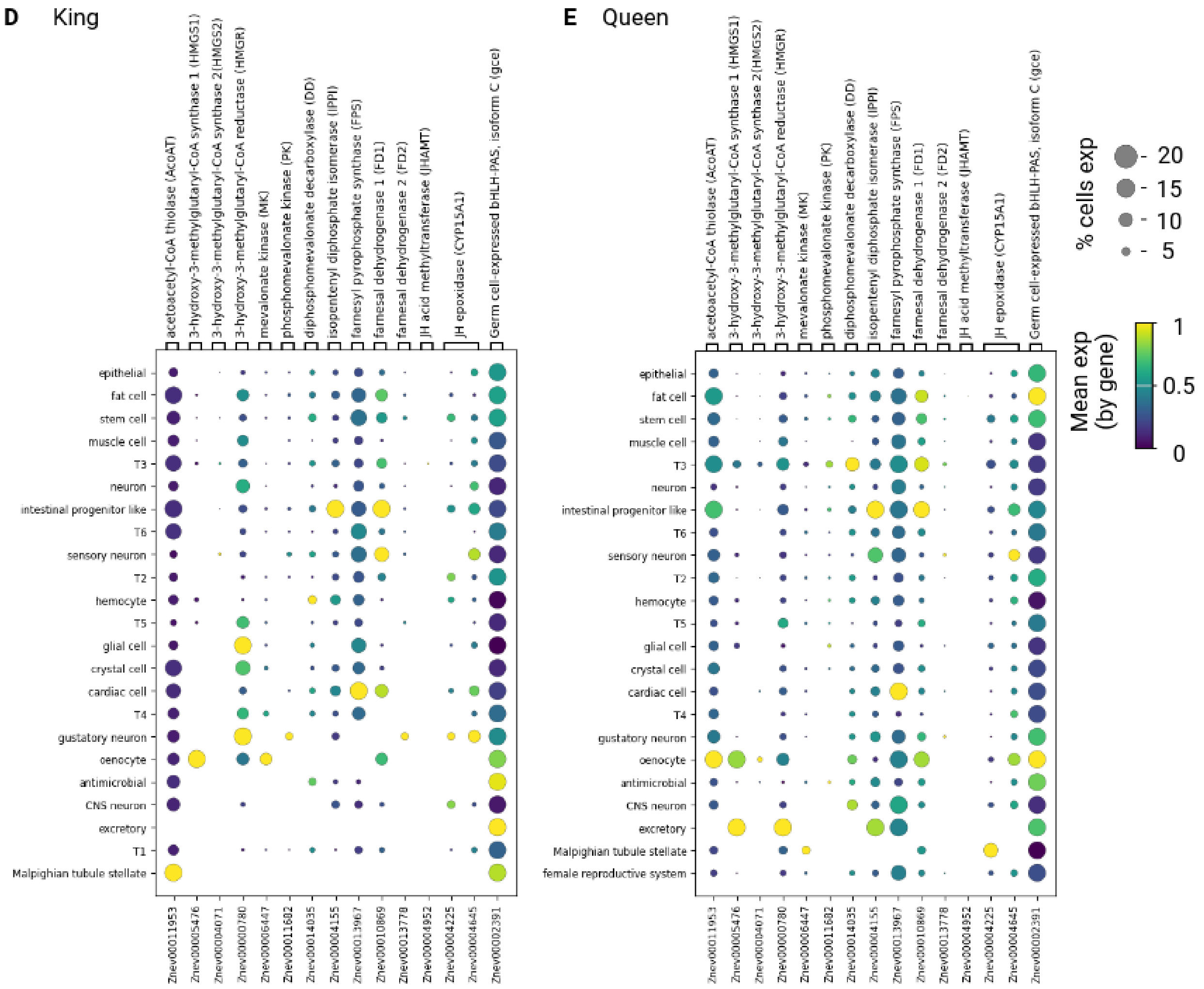
D
King
acetoacetyl-CoA thiolase (AcoAT)
3-hydroxy-3-methylglutaryl-CoA synthase 1 (HMGS1)
3-hydroxy-3-methylglutaryl-CoA synthase 2(HMGS2)
3-hydroxy-3-methylglutaryl-CoA reductase (HMGR)
mevalonate kinase (MK)
phosphomevalonate kinase (PK)
diphosphomevalonate decarboxylase (DD)
isopentenyl diphosphate isomerase (IPPI)
farnesyl pyrophosphate synthase (FPPS)
farnesal dehydrogenase 1 (FD1)
farnesal dehydrogenase 2 (FD2)
JH acid methyltransferase (JHAMT)
JH epoxidase (CYP15A1)
Germ cell-expressed bHLH-PAS, isoform C (gce)
epithelial
fat cell
stem cell
muscle cell
T3
neuron
intestinal progenitor like
T6
sensory neuron
T2
hemocyte
T5
glial cell
crystal cell
cardiac cell
T4
gustatory neuron
oenocyte
antimicrobial
CNS neuron
excretory
T1
Malpighian tubule stellate
Znev00011953
Znev00005476
Znev00004071
Znev00000780
Znev00006447
Znev00011602
Znev00014035
Znev00004155
Znev00013967
Znev00010869
Znev00013778
Znev00004952
Znev00004225
Znev00004645
Znev00002391
E
Queen
acetoacetyl-CoA thiolase (AcoAT)
3-hydroxy-3-methylglutaryl-CoA synthase 1 (HMGS1)
3-hydroxy-3-methylglutaryl-CoA synthase 2(HMGS2)
3-hydroxy-3-methylglutaryl-CoA reductase (HMGR)
mevalonate kinase (MK)
phosphomevalonate kinase (PK)
diphosphomevalonate decarboxylase (DD)
isopentenyl diphosphate isomerase (IPPI)
farnesyl pyrophosphate synthase (FPPS)
farnesal dehydrogenase 1 (FD1)
farnesal dehydrogenase 2 (FD2)
JH acid methyltransferase (JHAMT)
JH epoxidase (CYP15A1)
Germ cell-expressed bHLH-PAS, isoform C (gce)
epithelial
fat cell
stem cell
muscle cell
T3
neuron
intestinal progenitor like
T6
sensory neuron
T2
hemocyte
T5
glial cell
crystal cell
cardiac cell
T4
gustatory neuron
oenocyte
antimicrobial
CNS neuron
excretory
Malpighian tubule stellate
female reproductive system
Znev00011953
Znev00005476
Znev00004071
Znev00000780
Znev00006447
Znev00011602
Znev00014035
Znev00004155
Znev00013967
Znev00010869
Znev00013778
Znev00004952
Znev00004225
Znev00004645
Znev00002391
% cells exp
20
15
10
5
Mean exp
(by gene)
1
0.5
0

F

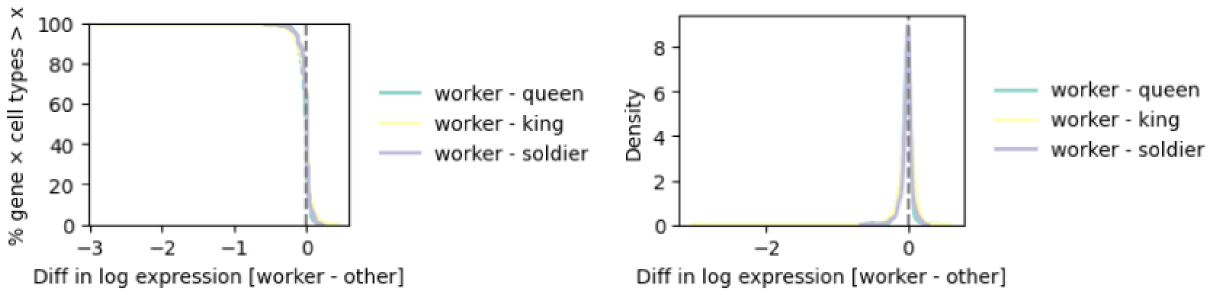


G

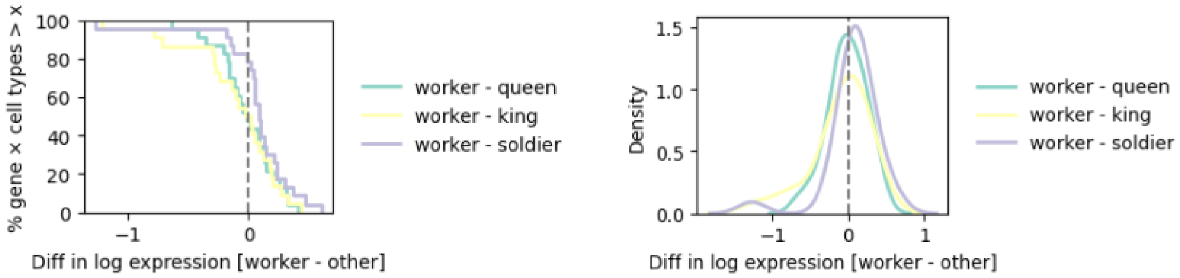


**Figure S5: Caste- and cell type-specific expression of developmental and functional genes. A)** Dot plots showing expression of the subset of juvenile hormone (*JH*) biosynthesis-related genes in the worker, soldier, king and queen across a subset of cell types (columns). **B) - D)** Dot plots showing expression of all of the juvenile hormone (*JH*) biosynthesis-related genes in the worker (**B**), soldier (**C**), king (**D**) and queen (**E**) across all cell types (columns). **F)** Complementary ECDF and kernel density plots of caste differences in expression across all genes in the juvenile hormone (JH) biosynthesis pathway. **G)** Same analysis as in (**H**), restricted to the JH receptor gene *gce*.

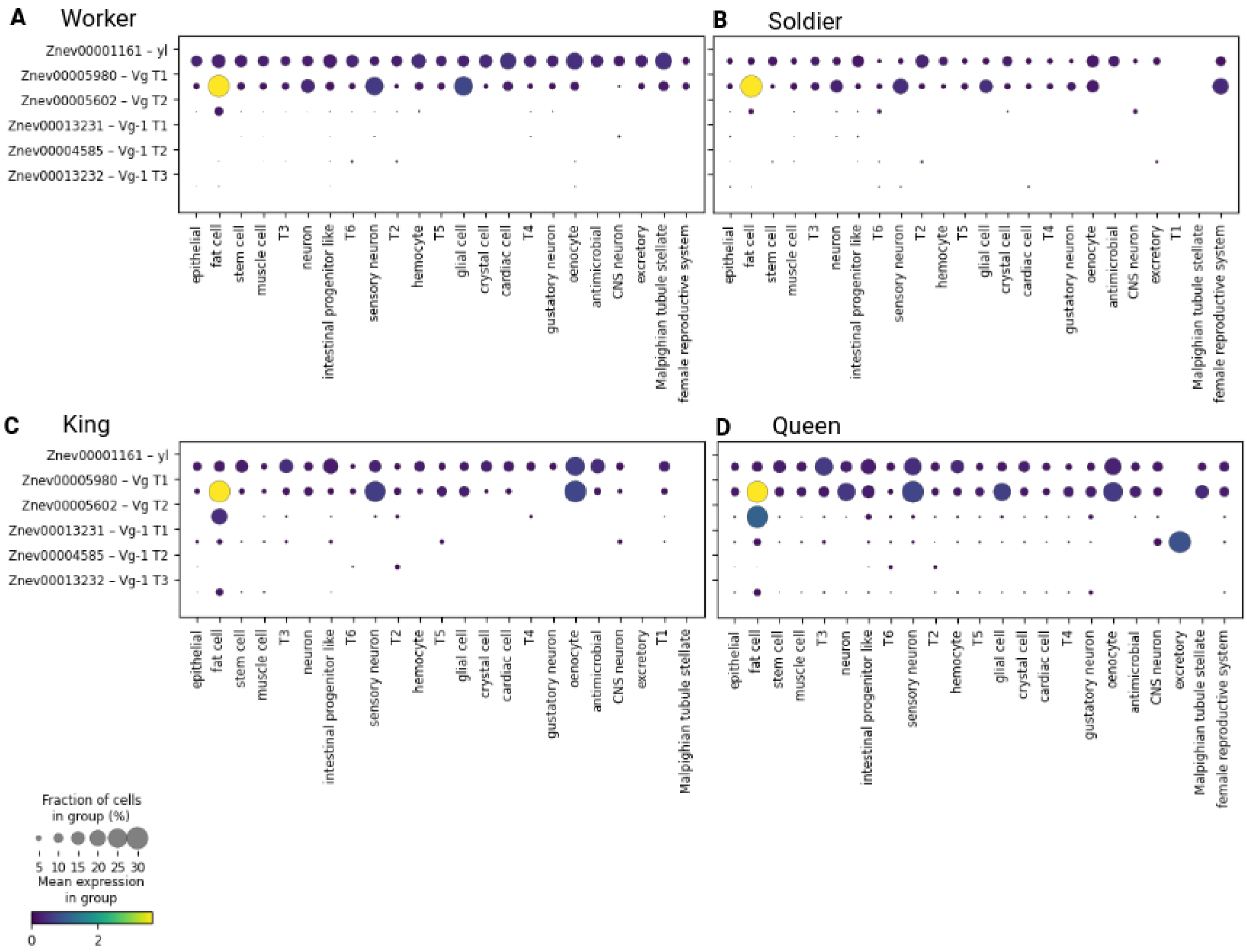


**Figure S6: Vitellogenin (Vg)-related genes across all castes. A-D)** Dot plots showing expression of Vitellogenin (*Vg*)-related genes across all cell types (columns) and castes (rectangles). The legend is shared between all panels. **A)** Expression in worker caste. **B)** Expression in soldier caste. **C)** Expression in king caste. **D)** Expression in queen caste.

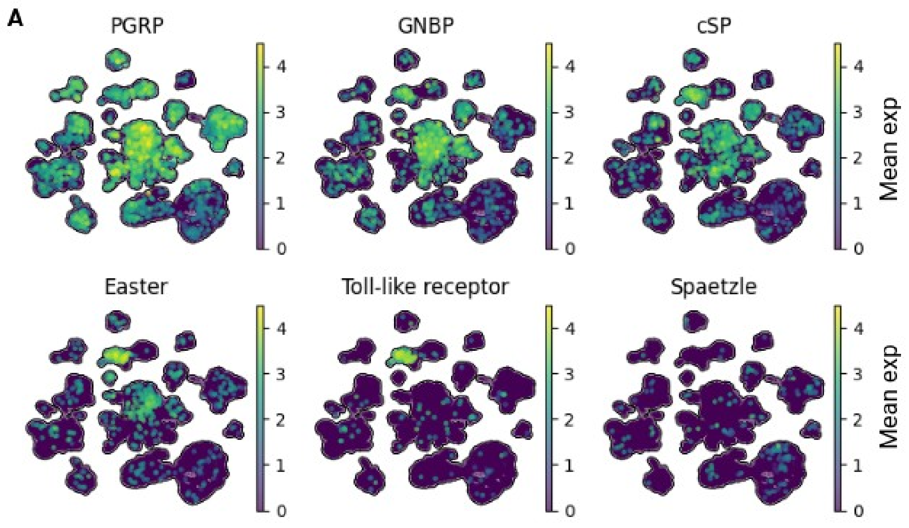


**Figure S7: Toll Signalling Cascade. A)** UMAP embedding of all nuclei coloured by expression of genes involved in the toll signalling cascade.

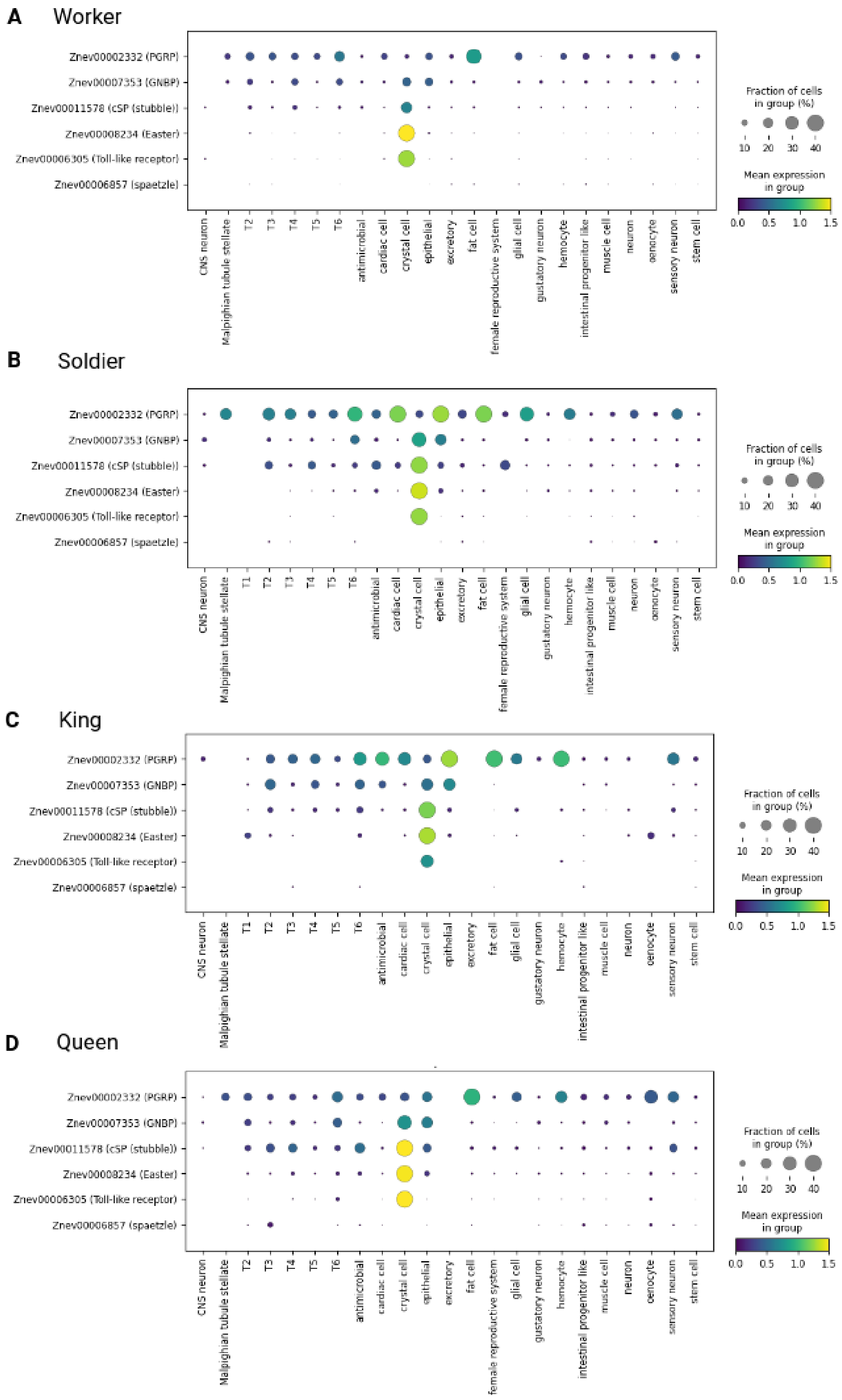

A
Worker
B
Soldier
C
King
D
Queen
Znev00002332 (PGRP)
Znev00007353 (GNBP)
Znev00011578 (cSP (stubble))
Znev00008234 (Easter)
Znev00006305 (Toll-like receptor)
Znev00006857 (spaetzle)
CNS neuron
Malpighian tubule stellate
T1
T2
T3
T4
T5
T6
antimicrobial
cardiac cell
crystal cell
epithelial
excretory
fat cell
female reproductive system
glial cell
gustatory neuron
hemocyte
intestinal progenitor like
muscle cell
neuron
oenocyte
sensory neuron
stem cell
Fraction of cells in group (%)
10 20 30 40
Mean expression in group
0.0 0.5 1.0 1.5

**Figure S8: Toll Signalling Cascade across all castes and cell types. A-D)** Dot plots showing expression of toll signalling cascade-related genes across all cell types (columns) and castes (rectangles). The legend is shared between all panels. **A)** Expression in worker caste. **B)** Expression in soldier caste. **C)** Expression in king caste. **D)**Expression in queen caste.

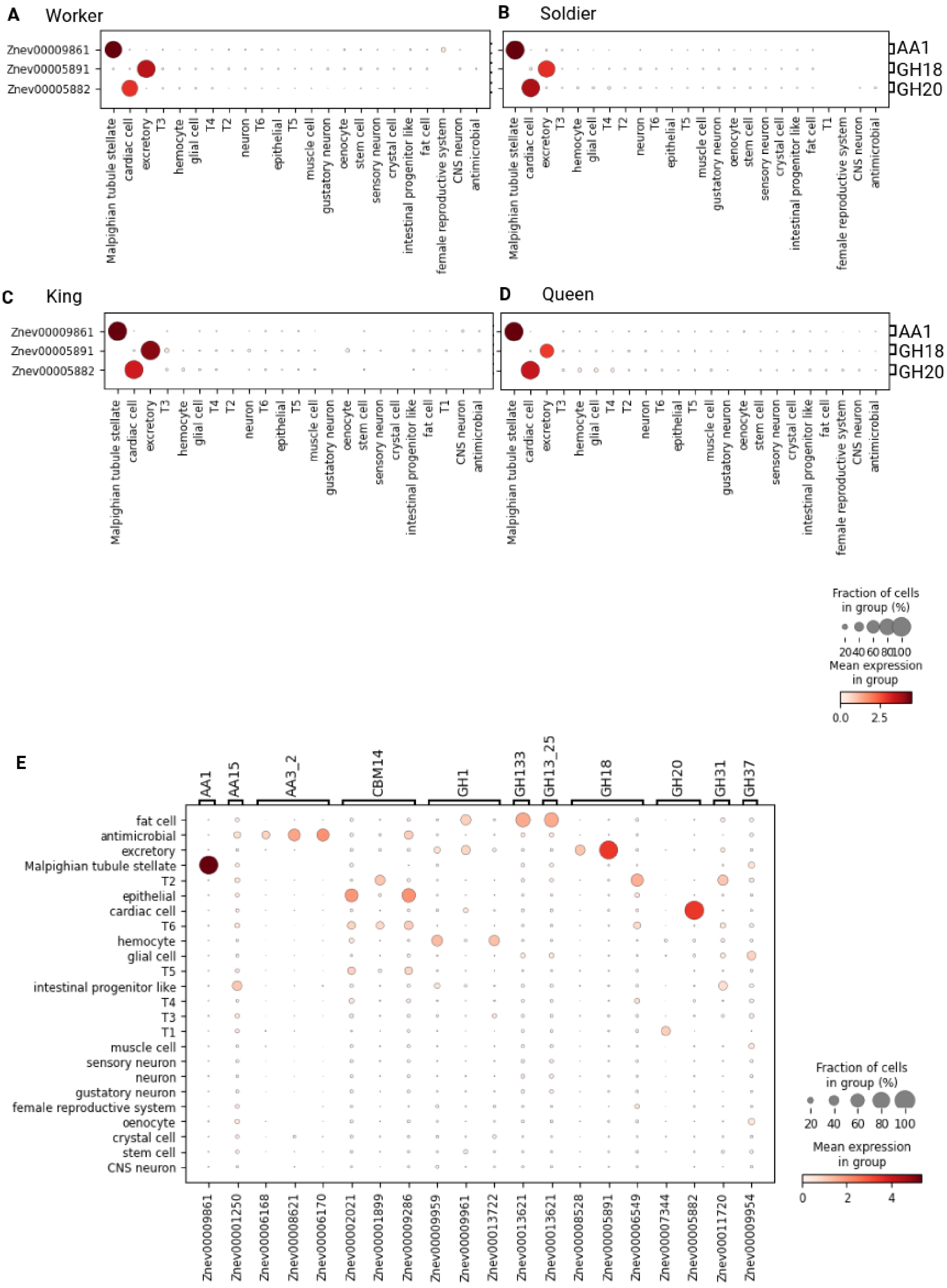
A Worker
B Soldier
C King
D Queen
Znev00009861
Znev00005891
Znev00005882
AA1
GH18
GH20
Malpighian tubule stellate
cardiac cell
excretory
T3
hemocyte
glial cell
T4
T2
neuron
T6
epithelial
T5
muscle cell
gustatory neuron
oenocyte
stem cell
sensory neuron
crystal cell
intestinal progenitor like
fat cell
T1
female reproductive system
CNS neuron
antimicrobial
Fraction of cells in group (%)
20 40 60 80 100
Mean expression in group
0.0 2.5
E
AA1
AA15
AA3_2
CBM14
GH1
GH133
GH13_25
GH18
GH20
GH31
GH37
Znev00009861
Znev00001250
Znev00006168
Znev00008621
Znev00006170
Znev00002021
Znev00001899
Znev00009286
Znev00009959
Znev00009961
Znev00013722
Znev00013621
Znev00013621
Znev00008528
Znev00005891
Znev00006549
Znev00007344
Znev00005882
Znev00011720
Znev00009954
Fraction of cells in group (%)
20 40 60 80 100
Mean expression in group
0 2 4

**Figure S9: CAZyme across all castes and cell types. A-D)** Dot plots showing expression of top three highest expressed CAZymes across all cell types (columns) and castes (rectangles). The legend is shared between all panels. **A)** Expression in worker caste. **B)** Expression in soldier caste. **C)** Expression in king caste. **D)**Expression in queen caste. **E)** Dot plots showing expression of top twenty highest expressed CAZymes across all cell types.

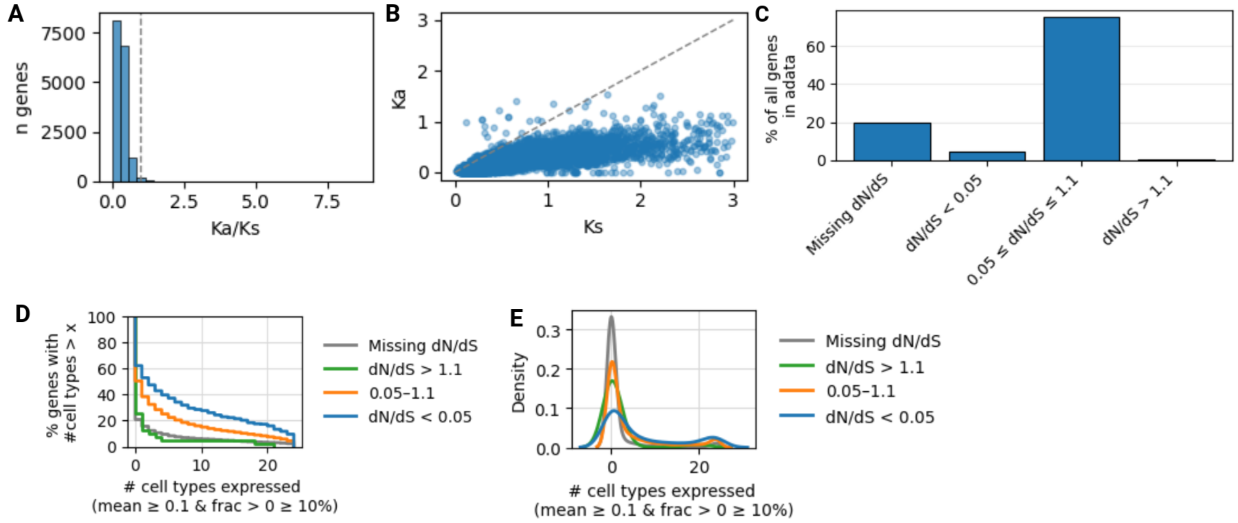


**Figure S10: Ka/Ks Values. A)** Distribution of nonsynonymous to synonymous substitution ratios (Ka/Ks) for genes present. The dashed vertical line indicates Ka/Ks = 1, corresponding to the neutral expectation. **B)** Scatter plot of nonsynonymous substitution rates (Ka) versus synonymous substitution rates (Ks) for orthologous genes between *H. sjostedti* and *Z. nevadensis.* Each point represents one gene. The dashed diagonal line indicates Ka = Ks. **C)** Bar chart of the proportion of genes grouped by dN/dS category (low (< 0.05, blue), middle (0.05-1.1, orange) and high (> 1.1, green)). **D)** Cumulative distribution of the number of cell types in which genes are expressed (mean log expression ≥ 0.1 in ≥ 10% of cells), grouped by low (< 0.05, blue), middle (0.05-1.1, orange) and high (> 1.1, green) Ka/Ks values. **E)** Density of the number of cell types in which genes are expressed (mean log expression ≥ 0.1 in ≥ 10% of cells), grouped by low (< 0.05, blue), middle (0.05-1.1, orange) and high (> 1.1, green) Ka/Ks values.
